# IL-27 induces a cytotoxic state in CD4^+^ T cells distinct from the Th1 lineage

**DOI:** 10.64898/2026.09.08.746061

**Authors:** Maria I. Matias, Remi Marrocco, Kyle Magro, Leonardo D. Sanchez Solis, Sabrina Figueroa Buezo, Laura Hinojosa-Gonzalez, Meronia Jabou, Haolin Lu, Erik Ehinger, Gabriel Ascui, Ye Zheng, Ferhat Ay, Chris A. Benedict, Samuel A. Myers

**Affiliations:** Laboratory for Immunochemical Circuits, La Jolla Institute for Immunology, La Jolla, CA, USA; Center for Autoimmunity and Inflammation, La Jolla Institute for Immunology, La Jolla, CA, USA; Division of Signaling and Gene Expression, La Jolla Institute for Immunology, La Jolla, CA, USA; Center for Cancer Immunotherapy, La Jolla Institute for Immunology, La Jolla, CA, USA; NOMIS Center for Immunology and Microbial Pathogenesis, Salk Institute for Biological Studies, La Jolla, CA, USA; Department of Pediatrics, University of California, San Diego, La Jolla, CA, USA; Center for Vaccine Innovation, La Jolla Institute for Immunology, La Jolla, CA, USA; Department of Pharmacology, University of California, San Diego, La Jolla, CA, USA; Program in Immunology, University of California, San Diego, La Jolla, CA, USA; Moores Cancer Center, UCSD Health, La Jolla, CA, USA

## Abstract

CD4^+^ cytotoxic T lymphocytes (CD4-CTLs) are understudied immune mediators with ambiguous origins. Despite expressing RUNX3, granzyme B (GZMB), and perforin (PRF1), CD4-CTLs are frequently classified as Th1 extensions due to shared interferon-γ (IFNγ) and T-BET expression. Here, we identify interleukin-27 (IL-27) as a independent inducer of a distinct CD4-CTL program. Proteomics reveals that while IL-27-polarized CD4+ T cells share protein signatures with conventional Th1s and CD8^+^ T cells, they possess a unique molecular landscape with re-wired cytokine signaling networks and a potent cytotoxic protein profile. Mechanistically, this program requires STAT1 and T-BET but operates independently of the autocrine IFNγ feedback that sustains Th1 cells. During acute murine cytomegalovirus infection, IL-27 receptor signaling contributes to CD4-CTL differentiation *in vivo*, as its loss leads to reduced GZMB expression and skews CD4^+^ T cells toward IFNγ^+^ and FOXP3^+^ subsets. Together, these findings establish IL-27 as a potent and previously unappreciated inducer of CD4-CTLs.

## Introduction

Cytotoxicity in the adaptive immune system is most widely attributed to CD8^+^ T cells, whereas CD4^+^ T cells conventionally coordinate immunity through distinct helper functions. However, CD4^+^ T cells can also acquire direct cytolytic capacity characterized by the expression of granzyme B (GZMB) and perforin (PRF1)^1,2^. While CD4-CTLs prominently emerge within the small intestinal CD4^+^ intraepithelial T cell (CD4-IET) compartment to maintain homeostasis^3,4^, they can also accumulate during viral infections and vaccinations, asthma, autoimmune diseases, and cancer^2^. Although CD4-CTLs are now recognized as a specialized subset within the CD4^+^ T cell compartment ^5^, the molecular mechanisms of their origin remain elusive. Despite being reported to emerge from various helper backgrounds ^2^, their differentiation is most frequently reported as a terminal differentiation of the Th1 subset^2,6–8^. This view stems from the frequent co-expression of GZMB with the canonical Th1 markers interferon-γ (IFNγ) and T-BET^9–12^, a phenotypic overlap that may obscure distinctions between Th1 cells and other cytotoxic CD4^+^ T cell subsets.

In the murine small intestine, a key step in cytotoxic CD4-IET generation is the loss of *Zbtb7b* (*Thpok*), the defining transcription factor for the CD4^+^ helper lineage^3,4^. This downregulation occurs alongside the upregulation of TBX21 (T-BET), and the CD8^+^ T cell cytotoxicity regulator RUNX3^13^, both of which are required for cytotoxic CD4-IET differentiation^3,4,14^. An *in vitro* model that mimics *in vivo* CD4-IETs was established using a combination of IL-27, TGFβ1, and retinoic acid (RA)^14^. TGFβ1, alone or in combination with RA, canonically drives the differentiation of FOXP3^+^ regulatory T cells (Tregs)^4,15,16^. Gut-adapted colonic Tregs can acquire a cytotoxic phenotype, including GZMB expression, to mediate tissue homeostasis, though these populations are rare^17^. IL-27, TGFβ1, and RA are constitutively present in the intestinal mucosa, yet the mechanism by which each signal integrates to divert CD4^+^ T cells from a common Treg state toward a CD4-IET-like cytotoxic fate remains poorly understood.

IL-27 is a heterodimeric cytokine composed of IL27 (p28) and IL27B (EBI3) subunits that signals through IL27RA (a.k.a. WSX-1) and the common co-receptor subunit IL6ST (a.k.a. GP130). IL-27 is a multifunctional, context-specific cytokine that induces both pro- and anti-inflammatory responses across a range of conditions^18,19^. In both CD8^+^ T cells and NK cells, IL-27 promotes cytotoxic function^20–22^. Within the CD4^+^ T cell compartment, IL-27 is classically recognized for its capacity to promote Type 1 regulatory (Tr1) cell differentiation, defined by the secretion of the anti-inflammatory cytokine IL-10^23–25^. However, emerging evidence suggests that IL-27 also serves as a potent driver of CD4^+^ T cell effector programs^18^. For instance, IL-27 signaling has been shown to upregulate critical activation and co-inhibitory receptors^26^ and drive pathogenic Th1 effector responses during autoimmune inflammation^27^. In the context of murine cytomegalovirus (MCMV) infection, IL-27 is required for the expansion of the IL-10^+^ CD4^+^ T cell pool which delay viral clearance^28^.

Given its established capacity to induce cytotoxicity in other CD8 and NK lineages^20–22^ and its capacity to promote CD4^+^ T cell effector responses^18^, we investigated whether IL-27 directly drives CD4-CTL differentiation as a distinct phenotypic fate rather than an extension of the conventional Th1 response. Uncoupling IL-27 from TGFβ1 and RA, we found that IL-27 alone acts as a potent driver of GZMB and PRF1 expression in CD4^+^ T cells, generating a more robust cytotoxic state than established IET-like CD4-CTL populations. Proteomic profiling shows that despite sharing core transcriptional regulators with Th1 cells (T-BET and IFNγ), IL-27 drives a molecular program that acquires CD8^+^ T cell cytotoxic features while remaining proteomically distinct from both Th1 and CD8^+^ T cells. Whereas Th1 cell maintenance depends on an autocrine IFNγ feedback loop^29,30^, this IL-27-driven cytotoxic program persists independently of such feedback. Functionally, IL-27-polarized CD4^+^ T cells rely on contact-dependent cytotoxicity, enabling chimeric antigen receptor (CAR)-expressing CD4^+^ T cells to specifically lyse cancer cells *in vitro*. Mechanistically, this IL-27-driven trajectory requires STAT1 to induce T-BET and RUNX3, with T-BET primarily contributing to GZMB expression. Using an *Il27ra* conditional knockout model during MCMV infection, we show that IL-27 signaling promotes differentiation of antigen-specific cytotoxic CD4^+^ T cells *in vivo*. Together, these findings uncover a previously unrecognized role of IL-27 in CD4^+^ T cells, demonstrating that rather than amplifying canonical Th1 responses, IL-27 acts as a standalone inducer of a CD4^+^ T cell cytotoxic program.

## Results

### IL-27 acts as an independent inducer of the CD4 cytotoxic program

CD4-IETs represent a specialized cytotoxic CD4 subset in the gut induced by the combination of signals from IL-27, TGFβ1, and RA^14^. However, the individual contribution of each signal in establishing this CD4-CTL program remains understudied. To evaluate how these cues direct cell fate, we established an *in vitro* differentiation model using naive CD4^+^ T cells from *Zbtb7b^GFP^ Runx3^tdTomato^*reporter mice, activated with α-CD3/α-CD28 agonist antibodies in the presence of IL-27 alone or in combination with TGFβ1 and RA for six days (gating strategy, **Extended Data Fig. 1a)**. Because TGFβ and RA are canonical drivers of Treg differentiation, induced Treg (iTreg) polarizing conditions (IL-2 + TGFβ ± RA) were included as reference controls for the regulatory phenotype, alongside unpolarized (Th0) controls (IL-2 + α-IFNγ + α-IL4) **(Fig. 1)**.

**Fig. 1.**
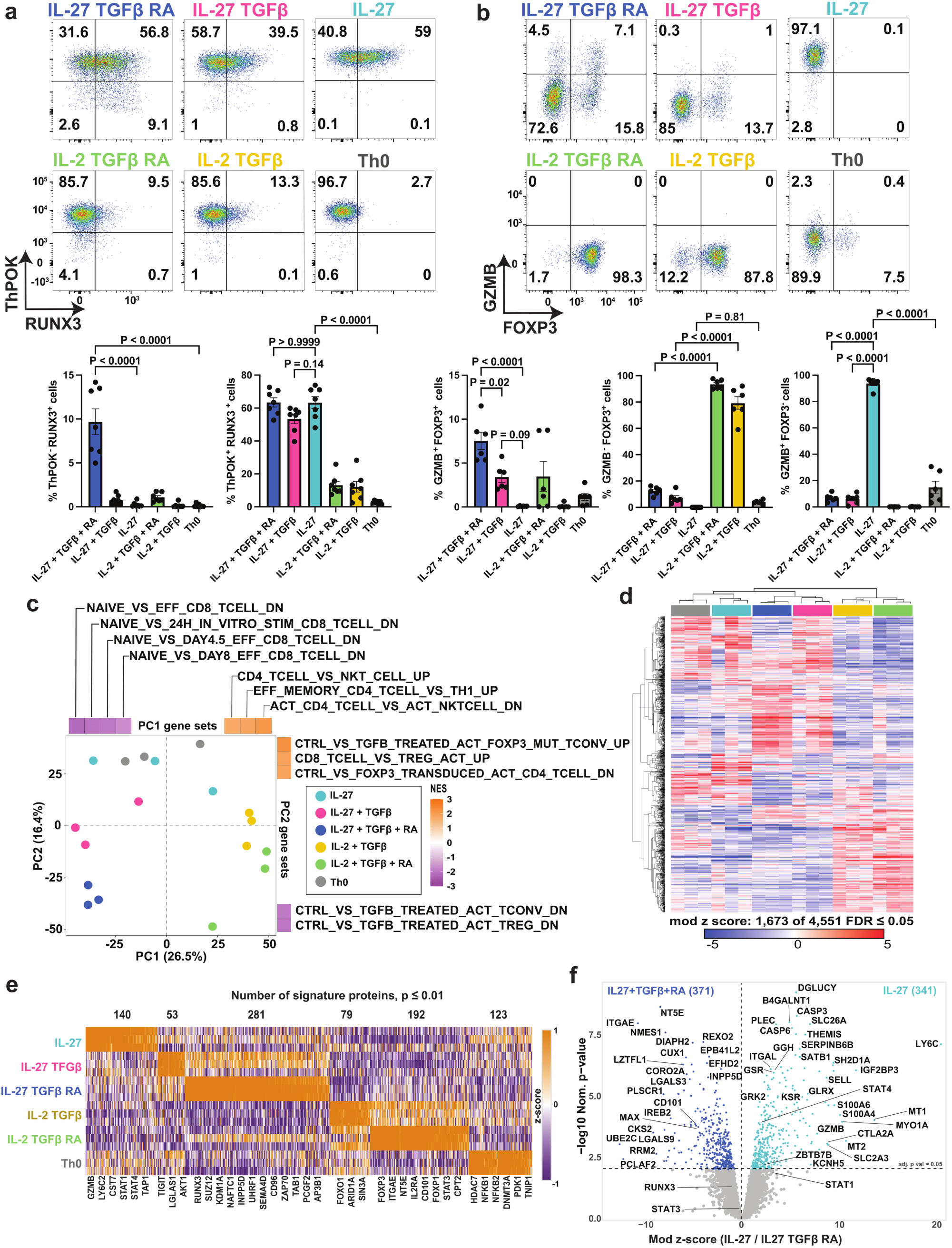
IL-27 acts as an independent inducer of the CD4 cytotoxic program. Naive CD4^+^ T cells from *Zbtb7b*^Gfp^ *Runx3*^tdTomato^ mice were activated with α-CD3 and α-CD28 agonist antibodies and polarized *in vitro* under the indicated conditions for 6 days. **a,b,** Representative flow cytometry plots (top) and summary quantitative bar graphs (bottom) detailing the frequencies of ThPOK^+^ (ZBTB7B) and RUNX3^+^ populations **(a)** and intracellular GZMB^+^ and/or FOXP3^+^ frequencies **(b)**. **c–f,** Global proteomic profiling of the polarized CD4^+^ T cells. **c,** Principal component analysis (PCA) of the proteomic data. Adjacent heatmaps indicate GSEA normalized enrichment scores (NES) of specific immunologic gene sets (MSigDB C7) correlating with PC1 and PC2. **d,** Unsupervised hierarchical clustering heatmap of differentially expressed proteins (moderated F-test) across the polarization conditions. The heatmap displays the relative abundance (row z-score) of 1,673 differentially abundant proteins (FDR ≤ 0.05) across the individual biological replicates for each polarization condition. **e,** Heatmap displaying the relative abundance (column z-score) of cell-specific proteins (marker selection analysis) across the varied polarization conditions. The total count of signature proteins defining each specific cluster is indicated above the heatmap. **f,** Volcano plot comparing differential protein abundance (moderated 2-sample T test) between the IL-27 + TGFβ + RA and IL-27 alone. Significantly enriched proteins are highlighted in navy (upregulated in IL-27 + TGFβ + RA) and teal (upregulated in IL-27). Proteins are labeled by their official gene symbol (e.g., ZBTB7B, TBX21) rather than common protein name (ThPOK, T-BET) throughout proteomics panels, consistent with Mouse Genome Informatics nomenclature. For **a** and **b**, data are presented as mean ± s.e.m. from n = 7 independent biological replicates; P values were determined by an ordinary one-way ANOVA with Tukey’s multiple comparisons test. For **c–f**, n = 3 independent mouse experiments per group; significance for marker proteins in **e** is defined as a nominal P ≤ 0.01, and the statistical threshold in f is set at an adjusted P value of ≤ 0.05.

We observed that the combination of IL-27, TGFβ1, and RA generated a distinct ThPOK^-^ RUNX3^+^ population (9.7%; **Fig. 1a, top left**), mimicking the phenotype of IET-like CD4-CTLs^3,4^ **(Fig. 1a)**. Removing RA abolished the terminal ThPOK^-^ RUNX3^+^ population and instead maintained an intermediate ThPOK^+^ RUNX3^+^ state (53%; **Fig. 1a, top middle**), consistent with the requirement for RA to drive terminal ThPOK downregulation. Notably, IL-27 administered separately from TGFβ1 and RA was sufficient to initiate expression of the cytotoxicity transcription factor RUNX3, yielding a prominent ThPOK^+^ RUNX3^+^ population (63%; **Fig. 1a, top right**). In contrast, iTreg and Th0 conditions maintained stable ThPOK levels with minimal induction of RUNX3.

Given that RUNX3 is a transcriptional driver of CD8^+^ T cell cytotoxicity^13,31^, we investigated GZMB induction across these diverse polarizing conditions. Because our model uses canonical Treg-inducing signals, we also monitored FOXP3 expression in parallel to track regulatory differentiation (**Fig. 1b**). The full IL-27 plus TGFβ and RA combination generated a cytotoxic GZMB^+^ FOXP3^-^ population (7%) alongside an intermediate GZMB^+^FOXP3^+^ double-positive fraction (8%) (**Fig. 1b, top left**). Omitting RA, leaving only IL-27 and TGF-β, severely repressed the cytolytic program, abrogating >85% of GZMB expression in both subsets (**Fig. 1b, top middle**). Unlike canonical iTreg conditions (IL-2 + TGFβ ± RA), which efficiently differentiated into classic FOXP3^+^ regulatory cells (79-93%), IL-27 + TGFβ ± RA conditions induced only modest FOXP3 upregulation (9-20%). Strikingly, IL-27 alone was sufficient to drive high GZMB expression (94%), establishing a prominent cytotoxic phenotype in the absence of FOXP3 (**Fig. 1b, top right**). Together, the phenotypes generated across these culture conditions span a continuum from GZMB^+^ cytotoxic IL-27-polarized cells to canonical FOXP3^+^ iTregs, bridged by double-positive intermediates.

To molecularly characterize this CD4^+^ T cell state continuum, we performed quantitative proteomic profiling of cells polarized for six days. Consistent with our observations by flow cytometry, mass spectrometry-based proteomic analysis confirmed a significant (P val = 0.002) inverse correlation between GZMB and FOXP3 protein levels across the six conditions **(Extended Data Fig. 1b)**. Principal component analysis (PCA) of the 4,551 proteins quantified revealed a bifurcation of the six conditions, with PC1 and PC2 capturing 42.6% of the cumulative variance **(Fig. 1c)**. PC1 (26.5% of variance) primarily delineated the cytokine treatments, separating IL-27-primed cells from IL-2. IL-27 alone clustered closely to Th0 on PC1 and PC2. Gene Set Enrichment Analysis (GSEA) of the proteins driving the separation in PC1 revealed an enrichment of terms associated with CD8^+^ T cell effectors **(Fig. 1c, Supplementary Table 1)**, demonstrating that IL-27 programs CD4^+^ T cells toward a cytotoxic profile that resembles *bona fide* cytotoxic CD8^+^ T cells. Conversely, conditions lacking IL-27 (IL-2 + TGFβ ± RA) retained conventional CD4^+^ T helper cell signatures but lacked cytotoxic and effector programs. GSEA of the PC2 loadings (16.4% variance) revealed that cells polarized with IL-27 alone retained a cytotoxic-associated signature. TGFβ1 addition drove a divergent shift along PC2, consistent with enrichment of TGF-β-dependent gene sets along that axis. Together, these data suggest that IL-27 alone establishes a core cytotoxic program, which is subsequently modified by TGFβ1 signaling.

To identify specific protein abundance differences between CD4^+^ T cell states, we performed a moderated F-test, identifying 1,673 differentially abundant proteins (FDR ≤ 0.05) in at least one polarization condition (**Fig. 1d** and **Supplementary Table 2**). Hierarchical clustering showed that both iTreg conditions were more similar to each other than to the other conditions, including Th0s. We next performed marker selection analysis to identify proteins relatively enriched in only one of the six CD4^+^ T cell polarization states **(Fig. 1e, Supplementary Table 3)**. We found that each CD4^+^ T cell polarization state had between 53 and 281 proteins that were increased in one cell state compared to all others (nominal p ≤ 0.01), where CD4-IET conditions had the largest signature. We identified a signature of 140 proteins upregulated by IL-27 alone that included GZMB, co-regulated modules associated with JAK/STAT signaling, Th1 polarization, and MHC-I antigen processing featuring STAT1, STAT4, and TAP1, as well as the surface marker LY6C, a previously reported response to IL-27 signaling^32^ **(Supplementary Table 3)**.

To isolate the proteomic footprint of each signal (IL-27, TGFβ1, or RA), we performed marker selection analysis (nominal p ≤ 0.01) by grouping conditions by the presence or absence of each factor (**Extended Data Fig. 1c–e**). This approach identified a core signature of 368 proteins consistently enriched across all IL-27-containing conditions **(Extended Data Fig. 1c)**. Conversely, tracking the total response to TGFβ1 isolated a shared signature of 329 proteins **(Extended Data Fig. 1d)**, while RA enriched a discrete module of 118 proteins (**Extended Data Fig. 1e**). Collectively, each stimulus maintains a discrete core proteomic profile regardless of co-applied stimuli.

Although IET-like CD4-CTLs served as our initial model, IL-27 alone drove stronger GZMB expression (**Fig. 1b**). To compare IL-27-polarized cells to this previously established cytotoxic model, we evaluated differential protein abundances between cells polarized with IL-27 alone versus IL-27 plus TGFβ and RA (**Fig. 1f**). As expected, GZMB was upregulated in IL-27 conditions alone, and CD4-IETs downregulated ZBTB7B (ThPOK). Adding TGFβ1 and RA to IL-27 significantly upregulated markers of tissue residency such as ITGAE (CD103), NT5E (CD73), and CD101. In contrast, cells polarized with IL-27 alone significantly upregulated the surface marker LY6C, the homing receptor SELL (CD62L), and the cytotoxic signaling adaptor SH2D1A. This direct comparison demonstrates that IL-27 alone drives an inflammatory cytotoxic program that is molecularly divergent from the tissue-resident features defining IET-like CD4-CTL. Collectively, these proteomic data demonstrate that IL-27 independently orchestrates a cytotoxic program in CD4^+^ T cells that is molecularly distinct from both unpolarized baselines and established IET-like CD4-CTL subsets.

### IL-27 drives a cytotoxic cellular state in CD4^+^ T cells distinct from the Th1 subset

CD4^+^ T cell cytotoxicity has been frequently attributed to the Th1 subset both *in vivo* and *in vitro*^2,33^. Given that our proteomic findings suggest IL-27 alone induces a molecular state overlapping with Th1 and *bona fide* cytotoxic CD8^+^ T cells (**Fig. 1c and 1e**), we directly compared these cell types. Naive CD4^+^ T cells from C57BL/6J mice were activated for three days under IL-27 alone or classically polarized Th1 conditions (IL-12 + α-IL4), alongside Th0 (IL-2 + α-IFNγ + α-IL4) cells as a neutral control and naive CD8^+^ T cells activated in presence of IL-2 as a positive cytotoxic control. Flow cytometry showed that IL-27-polarized cells were GZMB^+^ PRF^-^ to an extent similar to CD8^+^ T cells (58% and 67%, respectively), both significantly higher than Th1 cells (8%), and expressed PRF1 at slightly lower levels compared to CD8^+^ T cells (**Fig. 2a**). RUNX3 protein levels were highest in the IL-27 condition, exceeding both Th1 and CD8^+^ T cells (**Fig. 2b**).

**Fig. 2.**
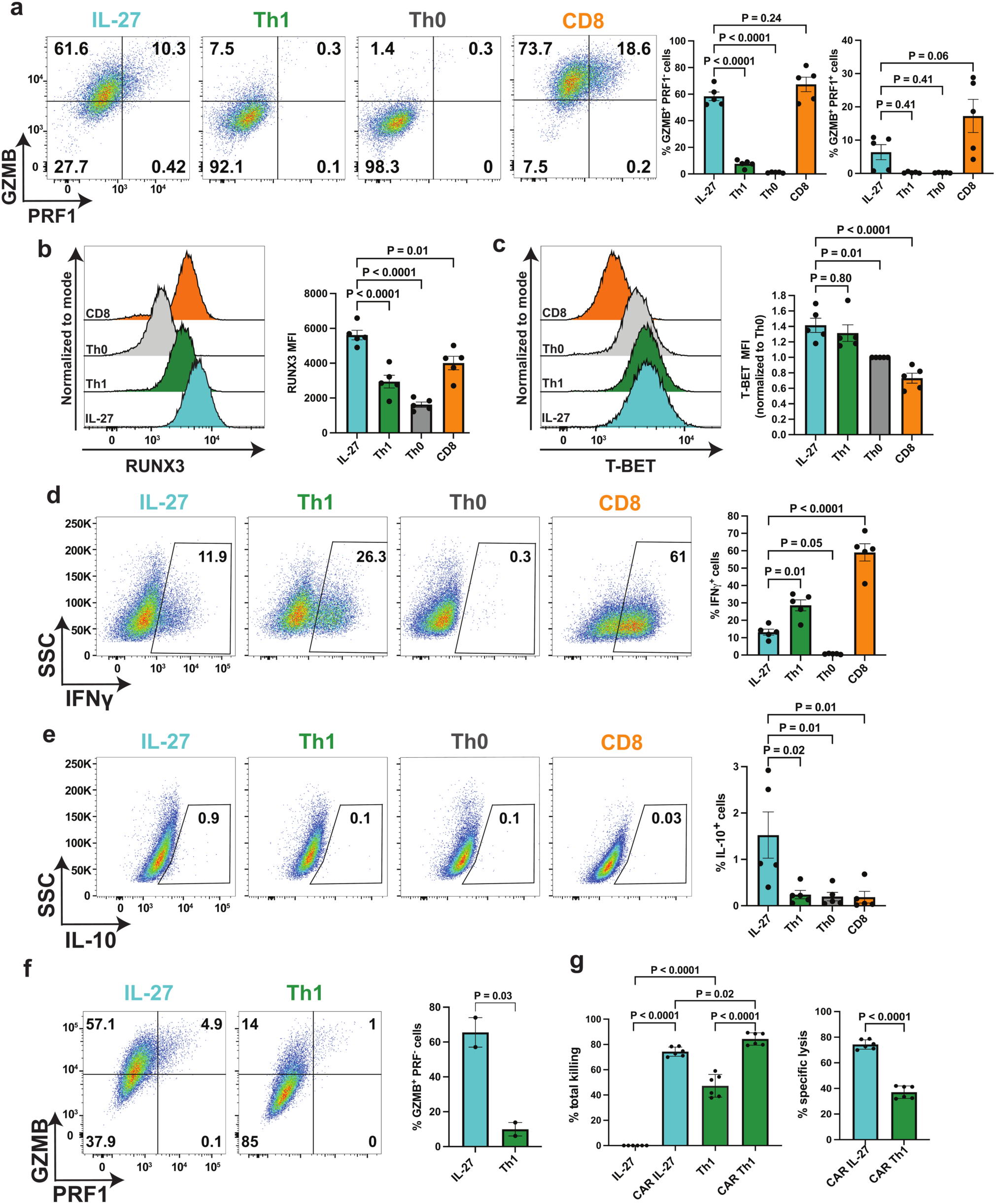
IL-27-polarized CD4^+^ T cells have overlapping maker gene expression with Th1s and CD8^+^ T cells. Naive CD4^+^ T cells from C57BL/6J mice were activated with α-CD3 and α-CD28 antibodies and polarized *in vitro* under the indicated conditions (IL-27, Th1, and Th0) for 3 days. Naive CD8^+^ T cells activated in the presence of IL-2 were included as positive cytotoxicity control. **a,** Representative flow cytometry plots (left) and summary bar graphs (right) detailing the frequencies of GZMB^+^ PRF1^−^ and GZMB^+^ PRF1^+^ populations. **b,c,** Representative flow cytometry histograms and summary bar graphs quantifying the mean fluorescence intensity (MFI) of RUNX3 **(b)** and T-BET **(c)**. T-BET MFI is normalized as fold-change relative to the Th0 condition. **d,e,** Representative flow cytometry plots and summary bar graphs assessing the frequencies of intracellular IFNγ^+^ **(d)** and IL-10^+^ **(e)** populations. **f,g,** *In vitro* cytotoxicity of engineered T cells. Mouse T cells were transduced with an anti-human CD19 chimeric antigen receptor (CAR) and differentiated under IL-27 or Th1 polarizing conditions for 3 days. **f,** Representative flow cytometry plots and summary bar graph showing frequencies of GZMB^+^ and PRF1^+^ populations in the CAR-transduced T cells prior to co-culture. **g,** CAR T cells were co-cultured with CD19^+^ Raji B target cells stably expressing luciferase for 48 hours. Bar graphs indicate the percentage of total target killing (left) and CAR-specific lysis (right) measured via luciferase activity loss. For **a–e**, data are presented as mean ± s.e.m. from n = 5 independent mouse replicates. For **f** and **g**, data are representative of 2 independent experiments, with **g** utilizing n = 3 technical replicates (3 independent wells) per condition, within each independent experiment. P values were determined by an ordinary one-way ANOVA with Tukey’s multiple comparisons test (**a–e**, and **g** left) or an unpaired two-tailed Student’s t-test (**f**, and **g** right).

Next, we examined subset marker expression between IL-27-polarized cells, Th1s, and CD8^+^ T cells. While both IL-27 and Th1 cells expressed similar levels of the CD4 subset marker T-BET (**Fig. 2c**), their cytokine profiles differed. Th1 cells, as expected, showed a high frequency of IFNγ-producing cells, whereas IL-27-induced cells showed a roughly 2.2-fold lower IFNγ^+^ frequency (**Fig. 2d**). Because IL-27 is one of several reported methods for producing IL-10^+^, FOXP3^-^ Tr1 cells^23–25^, we also assessed IL-10 production. IL-27-polarized cultures exhibited a statistically significant increase in the frequency of IL-10^+^ cells relative to Th1, Th0, and CD8s **(Fig. 2e),** and to levels consistent with previous reports^34^.

To assess whether IL-27 drives a cytotoxic state in human cells, primary human naive CD4^+^ T cells were activated for three days under the same polarizing conditions (**Extended Data Fig. 2**). IL-27 induced significantly higher frequencies of both GZMB^+^ PRF^-^ and GZMB^+^ PRF^+^ cells compared to Th0 controls. In contrast to the murine data, the frequency of single-positive GZMB^+^ PRF^-^ cells in human IL-27 conditions was comparable to Th1 cells (47% vs 51%). However, IL-27 treatment trended toward an increased frequency of double-positive GZMB^+^ PRF^+^ cells compared to Th1 conditions (16% vs 8%; **Extended Data Fig. 2a**). IL-27 drove a slight increase in T-BET compared to both Th1 and Th0 controls **(Extended Data Fig. 2b),** while RUNX3 expression was equivalent between IL-27 and Th1 cells **(Extended Data Fig. 2c)**. Consistent with the murine data, IL-27 induced lower levels of IFNγ compared to Th1 conditions **(Extended Data Fig. 2d)**. Collectively, these data demonstrate that IL-27 promotes a conserved cytolytic profile in primary both mouse and human CD4^+^ T cells.

To evaluate whether IL-27-polarized mouse CD4s were capable of antigen-specific killing, we performed an *in vitro* CAR-T killing assay (**Fig. 2f-g**). Primary naïve CD4^+^ T cells were activated for 18 hours and subsequently transduced to >80% efficiency with a human CD19-targeting CAR construct^35^ **(Extended Data Fig. 3a)**. Following transduction, the cells were cultured under either IL-27-supplemented or Th1-polarizing conditions. A 24-hour delay in IL-27 addition did not impact GZMB or PRF1 induction relative to cells exposed to IL-27 from the start of culture (**Fig. 2f)**. These engineered CD4^+^ CAR T cells were co-cultured with CD19^+^ Raji B target cells expressing luciferase at a fixed 10:1 effector-to-target (E:T) ratio, and target cell clearance was quantified after 48 hours of co-culture (**Fig. 2g**). As a baseline, Th1 cells exhibited high Raji cell death in the absence of the CAR, whereas IL-27-polarized cells showed undetectable basal killing (**Fig. 2g, left**). While both groups lysed target cells upon CAR expression, CAR-dependent cell killing was significantly higher in IL-27-CAR cells compared to Th1-CAR cells **(Fig. 2g, right)**, validating the contact-mediated cytotoxic function of these GZMB^+^ PRF1^+^ cells. Together, these results show that although IL-27-polarized CD4^+^ T cells share the subset-defining marker T-BET with Th1 cells, they diverge in cytokine output and exercise cytolytic function only upon antigen-specific engagement.

### IL-27-polarized CD4^+^ T cells are proteomically distinct from Th1s

Since IL-27-polarized CD4^+^ T cells share overlapping lineage markers with Th1s and CD8^+^ T cells, we used quantitative proteomics to characterize differences between these types after three days of *in vitro* differentiation. We first validated the T cell samples for proteomic analysis by comparing expression of key transcription factors (RUNX3 and T-BET) and the effector protein GZMB by flow cytometry (**Fig. 3a**). Expression patterns for these markers were concordant across both techniques, with IL-27-polarized cells showing the highest combined expression.

**Fig. 3.**
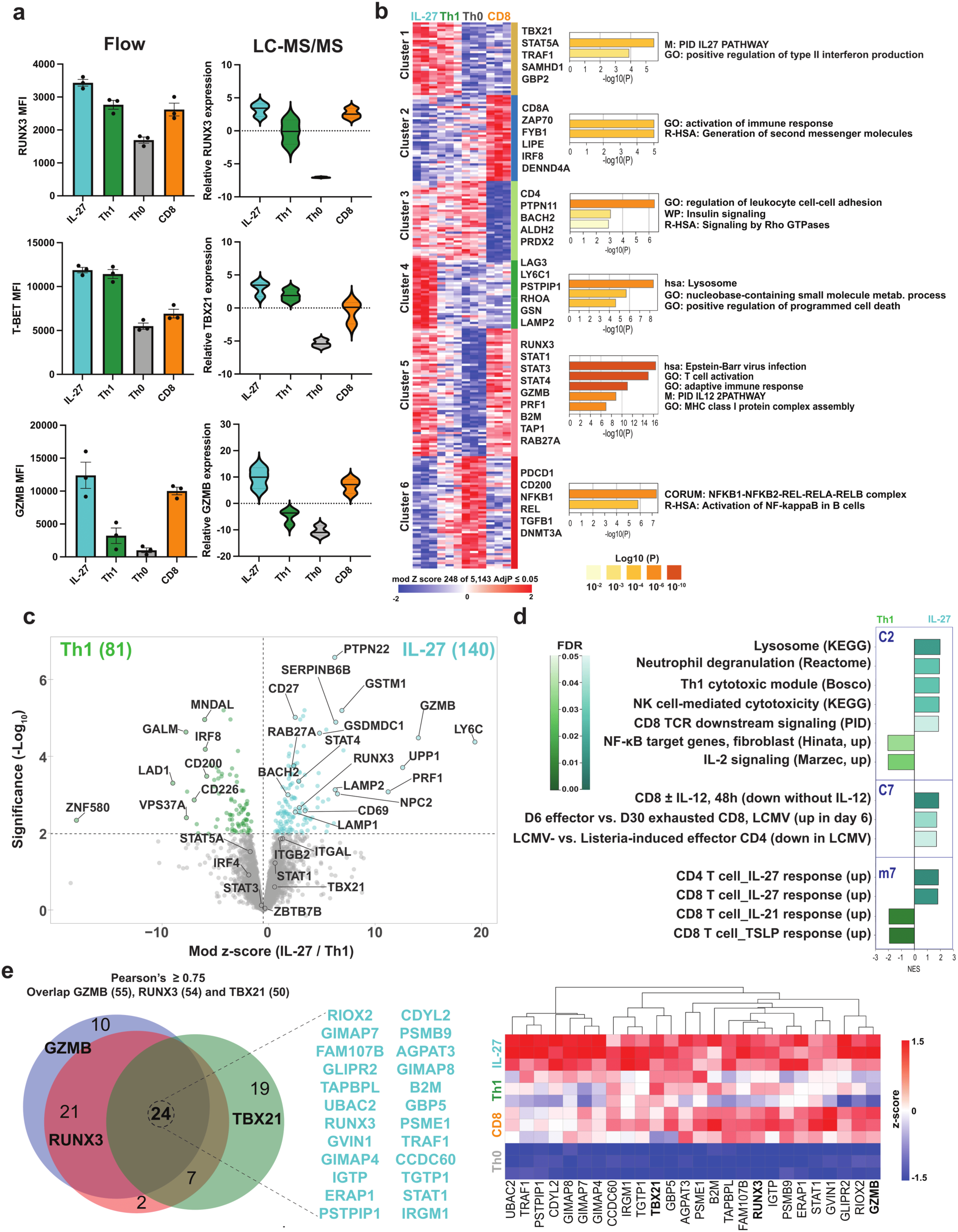
IL-27-polarized CD4^+^ T cells are proteomically distinct from the Th1 subset. Naive CD4^+^ T cells from *Zbtb7b*^Gfp^ *Runx3*^tdTomato^ mice were activated with α-CD3 and α-CD28 antibodies and polarized *in vitro* under the indicated conditions (IL-27, Th1, and Th0) for 3 days. Naive CD8^+^ T cells activated in the presence of IL-2 were included as positive cytotoxicity control. **a,** Cross-platform validation of core effector proteins and transcription factors. Bar graphs (left) denote the mean fluorescence intensity (MFI) of RUNX3, T-BET, and GZMB assessed via flow cytometry. Adjacent violin plots (right) display the corresponding relative protein expression levels captured by liquid chromatography-tandem mass spectrometry (LC-MS/MS). As in Fig. 1, gene symbols (e.g., TBX21) rather than its alias T-BET are used to label proteomics data throughout this figure. **b,** K-means clustering of the global proteomics (left) resolving distinct modules across the T cell states (Cluster 1 to 6). Row z-scores of the 248 differentially abundant proteins (moderated F-test, adjusted P ≤ 0.05) are displayed. Bar graphs (right) detail matched Gene Ontology (GO) pathway enrichment analysis for each respective cluster. **c,** Volcano plot detailing differential protein abundances between the IL-27 and Th1 CD4^+^ T cell conditions. Significantly enriched proteins are highlighted in teal (IL-27, 140 proteins) and green (Th1, 81 proteins). **d,** Gene Set Enrichment Analysis (GSEA) displaying Normalized Enrichment Scores (NES) of selected immunologic signatures (C2, C7, and m7 modules) enriched in the IL-27 versus Th1 cells. **e,** Protein correlation analysis defining the core co-expression module. Venn diagram (left) demonstrates the intersection of the differentially abundant proteins strongly correlated (Pearson’s correlation coefficient r ≥ 0.75) with the expression of GZMB, RUNX3, and TBX21. Heatmap (right) displays the relative abundance (column z-score) of the 24 uniquely intersecting core network proteins, alongside GZMB and TBX21, across the indicated T cell populations. For flow cytometry data in **a**, data are presented as mean ± s.e.m. from n = 3 independent biological replicates. For all proteomics data (**a** right, **b–e**), data are derived from n = 3 independent experiments.

To identify differential protein abundances, we performed a moderated F-test followed by K-means clustering to categorize differential protein abundance across the four T cell subsets (**Fig. 3b**, **Supplementary Table 4**). Six clusters were identified and annotated by Gene Ontology (GO) enrichment analysis to define their core biological functions. Cluster 1 was characterized by proteins shared between Th1 and IL-27-polarized cells (e.g., TBX21, STAT5A). This module was significantly enriched for the Pathway Interaction Database (PID) “IL-27 pathway term”, highlighting the reciprocal gene expression overlap between IL-27 polarization and Th1s. In contrast, cluster 4 was restricted to the IL-27 condition and enriched for lysosomal proteins, such as LAMP2, consistent with an active degranulation/lysosomal-trafficking phenotype. IL-27-polarized cells and CD8^+^ T cells shared cluster 5, containing core mediators including RUNX3, GZMB, and PRF1, alongside MHC class I components such as B2M and TAP1. This cluster was strongly enriched for GO terms including "MHC class I protein complex assembly" and the "PID IL12 pathway." Because IL-27-polarized cells and CD8^+^ T cells were differentiated without exogenous IL-12, this again suggests IL-27 independently drives gene expression that overlaps with IL-12 polarization.

To further delineate differences between IL-27-polarized cells and Th1s, which share overlapping subset markers (**Fig. 2 and 3a**) and protein signatures (**Fig. 3b**), we directly compared their proteomes. Pairwise comparison identified that while both subsets expressed comparable levels of STAT1 and TBX21 (T-BET), IL-27-polarized cells trended toward higher expression of ITGAL and ITGB2, the constituent subunits of the synapse-stabilizing integrin LFA-1. IL-27-polarized cells also showed significant upregulation of GZMB and PRF1, alongside the lytic granule trafficking machinery RAB27A, LAMP1 (CD107A), and LAMP2 (CD107B). IL-27 also significantly enriched for LY6C, RUNX3, and the costimulatory receptor CD27 **(Fig. 3c)**. GSEA analysis further confirmed this functional distinction, showing IL-27-polarized cells were significantly more enriched for the established ‘Th1 Cytotoxic Module’ than classical Th1 cells themselves **(Fig. 3d, Supplementary Table 5)**. In addition, IL-27-polarized cells showed positive enrichment for innate-like killing mechanisms, including NK cell-mediated cytotoxicity, lysosomal activity, and degranulation pathways, together with CD8^+^ TCR downstream signaling and functional effector T cell signatures, alongside negative enrichment for IL-2 and NF-kB signaling. Together, these data indicate that IL-27-polarized cells adopt a proteomic landscape that mirrors *bona fide* CD8^+^ and NK cell cytotoxicity, diverging from the IL-2 and NF-kB-driven activation program associated with conventional Th1 helper function.

Finally, we defined co-expression modules associated with the IL-27 cytolytic program using Pearson’s correlation analysis of all differential abundant proteins against GZMB, RUNX3, and TBX21 abundance (**Fig. 3e**, **Supplementary Table 6**). We selected these three markers as references because IL-27-polarized cells co-express all three at high levels, whereas Th1 cells show high T-BET with low RUNX3 and CD8^+^ T cells show high RUNX3 with intermediate T-BET. The intersection of these analyses revealed a signature of 24 proteins that significantly correlated with all three proteins (Pearson’s r ≥ 0.75) (**Supplementary Table 6)**. Hierarchical clustering showed these proteins were most enriched in IL-27-polarized and CD8^+^ T cells, reduced in Th1 cells, and lowest in Th0 controls (**Fig. 3e**). This 24-protein module includes the transcription factor STAT1 and a cluster of interferon-inducible GTPases (IRGM1, TGTP1, IGTP, GVIN1, and GBP5) that mediate cell-autonomous antimicrobial defense^36^. Pathway analysis indicated that this core is significantly enriched for antigen processing and presentation, the ER-phagosome pathway, and NOD-like receptor signaling **(Extended Data Fig. 4)**. Collectively, these results indicate that while IL-27 and Th1 cells share a core Type 1 identity, IL-27 drives a specialized program characterized by a cytolytic and structural protein signature shared with CD8^+^ CTLs.

### IL27-polarized cells have a re-wired cytokine signaling network compared to Th1 cells

In Th1 cells, IL-12 signals through STAT4, driving IFNγ expression and a positive feedback loop that reinforces both IFNγ and T-BET expression^29,30^. Given this reliance on autocrine IFNγ signaling in Th1 cells, we investigated whether the IL-27-driven program is similarly maintained by an autocrine IFNγ feedback loop. We activated naive CD4^+^ T cells in IL-27 or Th1 polarizing conditions in the presence or absence of α-IFNγ neutralizing antibodies **(Fig. 4)**. Blocking IFNγ significantly reduced IFNγ production in Th1 cells six-fold, while the percentage of IFNγ^+^ cells in the IL-27 condition was unaffected **(Fig. 4a)**. Consistent with the known Th1-promoting feedback loop, blocking IFNγ significantly reduced T-BET and RUNX3 levels in Th1 cells **(Fig. 4b, c).** IL-27-polarized cells maintained high expression of both transcription factors regardless of IFNγ blockade. GZMB and PRF1 levels were not affected by autocrine IFNγ signaling in either IL-27-polarized cells or Th1s **(Fig. 4d)**. Because IL-27 has also been shown to induce an autocrine IL-21 feedback loop that promotes cytotoxicity in CD8^+^ T cells^37,38^, we tested whether IL-21 blockade affected GZMB or PRF1 expression and found no effect in either condition **(Extended Data Fig. 5**). Collectively, these results demonstrate that IL-27-polarization sustains a CD4-CTL identity independent of IFNγ and IL-21.

**Fig. 4.**
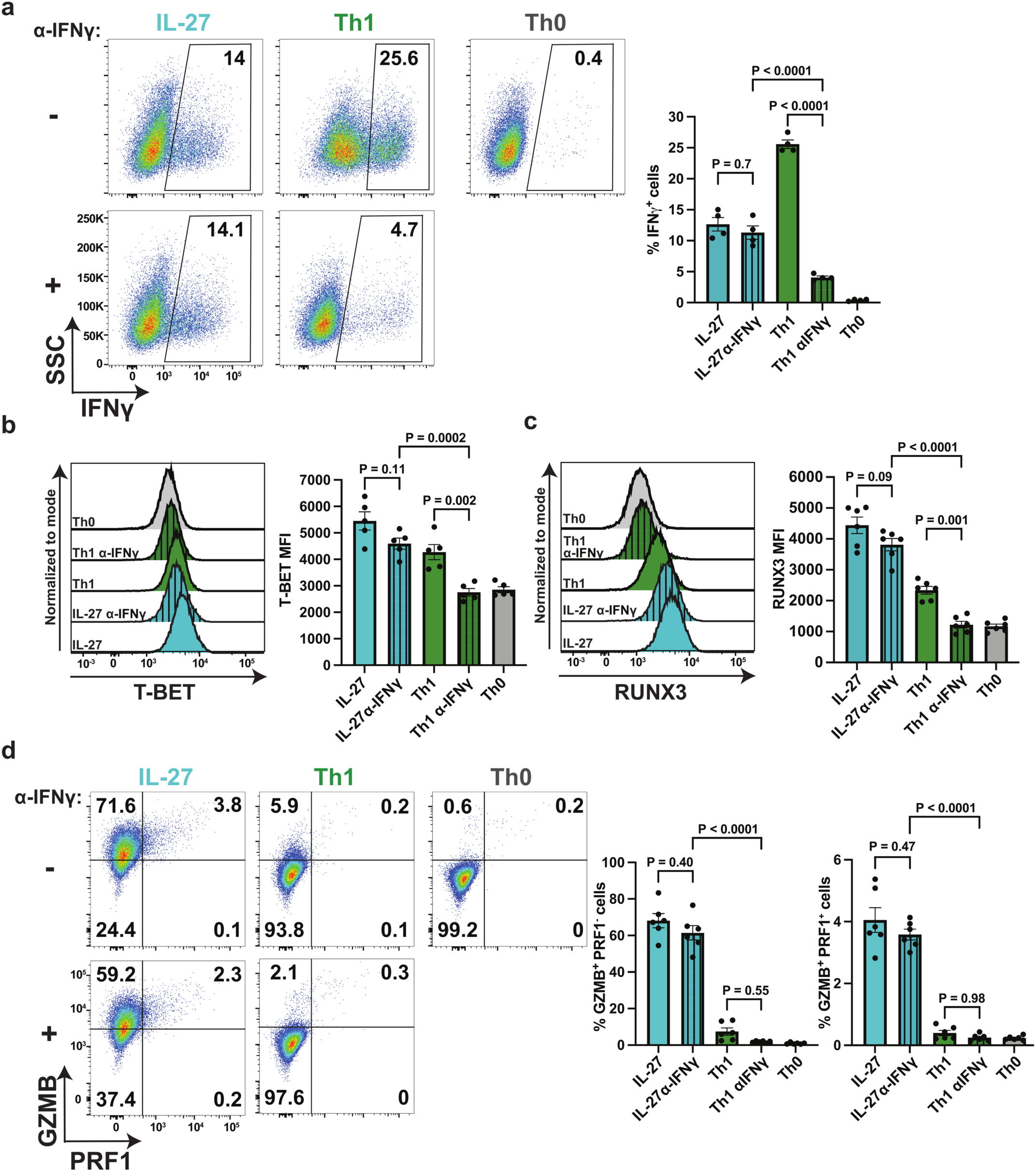
IL-27 drives CD4-CTL differentiation independently of the Th1-associated autocrine IFNγ feedback. Naive CD4^+^ T cells from C57BL/6J mice were activated with α-CD3 and α-CD28 antibodies and polarized *in vitro* under the indicated conditions (IL-27, Th1, and Th0) for 3 days in the presence (+) or absence (–) of a neutralizing antibody targeting IFNγ (α-IFNγ). **a,** Representative flow cytometry plots (left) and summary bar graph (right) assessing the frequency of intracellular IFNγ^+^ cells following α-IFNγ blockade. **b,c,** Representative flow cytometry histograms and summary bar graphs quantifying the mean fluorescence intensity (MFI) of T-BET **(b)** and RUNX3 **(c)** across the indicated conditions. **d,** Representative flow cytometry plots (left) and summary bar graphs (right) detailing the frequencies of GZMB^+^ PRF1^−^ and GZMB^+^ PRF1^+^ populations following α-IFNγ blockade. Data are presented as mean ± s.e.m. from n = 4–6 independent biological replicates. P values were determined by an ordinary one-way ANOVA with Tukey’s multiple comparisons test.

### STAT1 is required for the IL-27-mediated cytotoxic program

To investigate the transcriptional regulation of GZMB in IL-27 conditions, we examined STAT transcription factor activation. In T cells, IL-27 is known to signal through STAT1, STAT3, and STAT5^19^. To verify this signaling in our model, we activated naive CD4^+^ T cells for 30 minutes in the presence of no cytokines, IL-12, or IL-27, and stained for STAT phosphorylation. As expected, IL-27 induced phosphorylation of STAT1 Y701, STAT3 Y705, and STAT5 Y694 (**Fig. 5a**). STAT4 Y693 phosphorylation was absent in both conditions, since the IL-12 receptor subunit IL12RB2 is not expressed until about 16-24 hours into CD4^+^ T cell activation^39^. Given that our proteomics identified STAT1 as a highly correlated component of the IL-27-driven protein network, we tested its requirement for establishing the cytotoxic program by isolating naive CD4^+^ T cells from germline *Stat1* knockout (*Stat1^−/−^*) mice and evaluating their differentiation capacity. Loss of STAT1 significantly reduced the induction of RUNX3 and T-BET in both IL-27 and Th1 conditions, with no effect on the already-low RUNX3 expression in Th0 cells **(Fig. 5b, c)**. We then assessed the impact of *Stat1* loss on the cytotoxic effector program **(Fig. 5d-e)**. In the absence of STAT1, IL-27-polarized cells failed to express GZMB and PRF1, abolishing the cytotoxic phenotype (75% vs 0.8% GZMB^+^PRF1^-^ in WT vs *Stat1*^-/-^) **(Fig. 5d)**. While IL-27 is known to drive STAT3-dependent transcriptional programs^23^, pharmacological inhibition of STAT3 using STATTIC^40^ during *in vitro* IL-27 differentiation did not impact the induction of this program (**Extended Data Fig. 6**). Finally, STAT1 was essential for IFNγ production in the IL-27 condition **(Fig. 5e)**, whereas only a small IFNγ population persisted in *Stat1*^-/-^ Th1 cultures **(Fig. 5e)**, likely reflecting the reliance on STAT4 to promote IFNγ expression^29^. Together, these results identify STAT1 as a key transcription factor required for IL-27 to drive the CD4-CTL program.

**Fig. 5.**
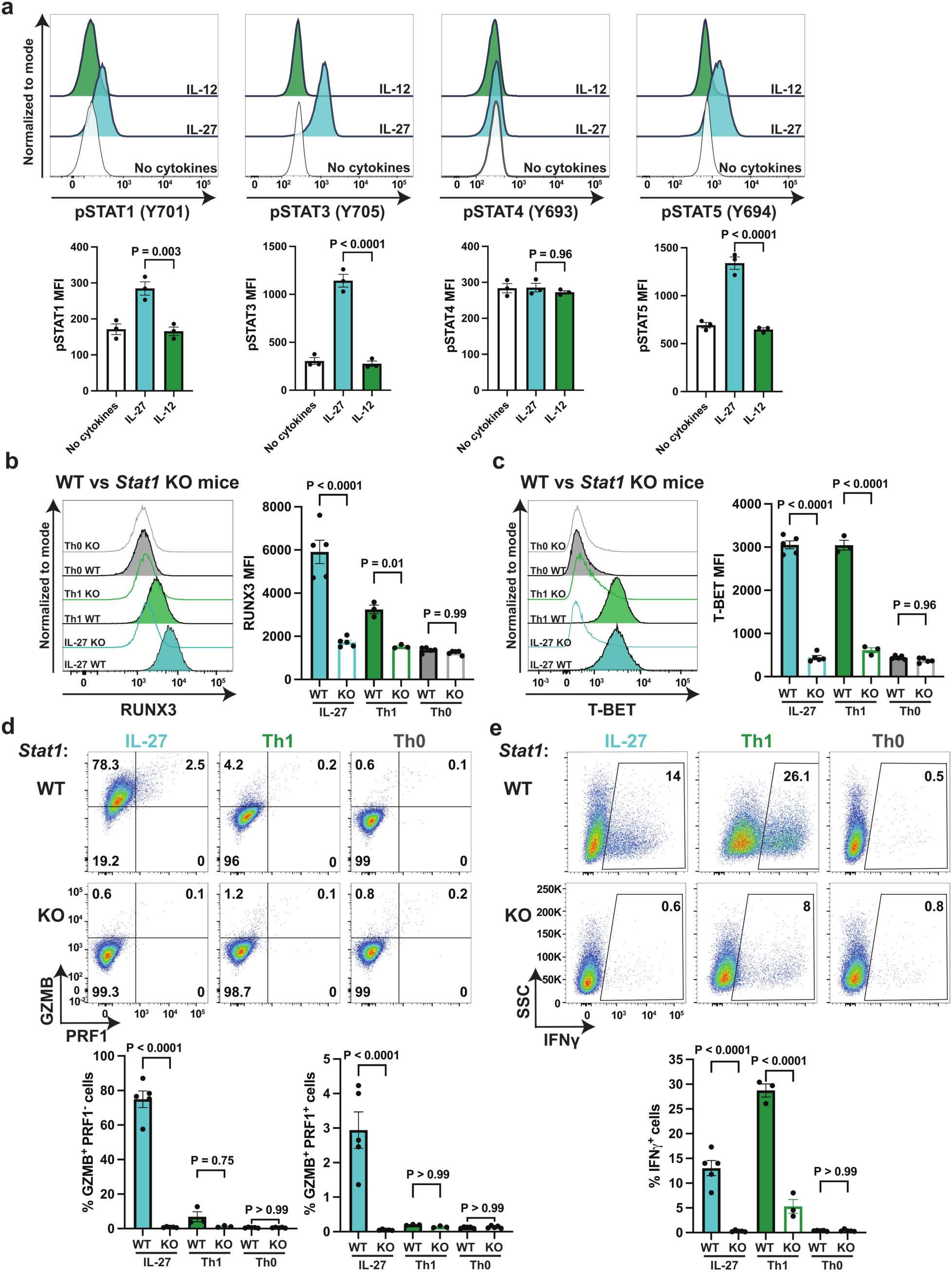
STAT1 is required for the IL-27-mediated cytotoxic program. **a,** PhosFlow cytometric analysis of STAT signaling. Naive CD4^+^ T cells from C57BL/6J mice were activated with α-CD3 and α-CD28 antibodies and stimulated with IL-27, IL-12, or left without cytokines for 30 minutes. Representative histograms (top) and summary bar graphs (bottom) depict the mean fluorescence intensity (MFI) of phosphorylated STAT1 (pSTAT1) Y701, pSTAT3 Y705, pSTAT4 Y693, and pSTAT5 Y694. **b–e,** Flow cytometric evaluation of cytotoxic differentiation in wild-type (WT) versus *Stat1*-knockout (KO) CD4^+^ T cells. Naive CD4^+^ T cells from C57BL/6J WT or *Stat1* KO mice were activated with α-CD3 and α-CD28 antibodies and polarized *in vitro* under the indicated conditions (IL-27, Th1, and Th0) for 3 days. **b,c,** Representative flow cytometry histograms and summary bar graphs quantifying the MFI of RUNX3 **(b)** and T-BET **(c)** across genotypes and polarization conditions. **d,** Representative flow cytometry plots (top) and summary bar graphs (bottom) detailing the frequencies of GZMB^+^ PRF1^−^ and GZMB^+^ PRF1^+^ populations. **e,** Representative flow cytometry plots (top) and summary bar graph (bottom) assessing intracellular IFNγ^+^ population frequencies. Data are presented as mean ± s.e.m. from n = 3– 6 independent biological replicates. P values were determined by an ordinary one-way ANOVA with Tukey’s multiple comparisons test.

### T-BET is essential for establishing the IL-27-driven cytotoxic program

To investigate the individual contributions of the regulators T-BET and RUNX3, we used Cas9-expressing mice to perform CRISPR-mediated deletion of these two transcription factors. Naive CD4^+^ T cells were activated and transduced with the respective sgRNAs 24 hours post-activation. Following transduction, cells were cultured for an additional 24 hours to allow for gene deletion prior to the addition of IL-27. Flow cytometric analysis showed a specific and significant reduction in protein levels for both T-BET and RUNX3 in cells transduced with their respective guide RNAs **(Fig. 6a)**. In this acute deletion model, loss of either transcription factor did not significantly impact the expression of the other (**Fig. 6a**). Because IL-27 was added 48 hours after initial activation, baseline GZMB levels in the non-targeting control (NTC) were ∼50% lower than in cells polarized immediately (**Fig. 6b**), consistent with reports that IL-27 timing and starting differentiation state influence the resulting phenotype^41^. Despite this, acute deletion of *Tbx21* significantly reduced both the frequency of GZMB^+^ cells and MFI, whereas *Runx3* deletion had a more modest and non-significant effect **(Fig. 6b)**. To corroborate these CRISPR-Cas9 results, we used germline *Tbx21* KO mice to assess the program under total T-BET deficiency (**Fig. 6c–e**). In *Tbx21* KO cells, RUNX3 induction was reduced 1.8-fold but not completely abolished compared to control C57BL/6J CD4^+^ T cells, suggesting a parallel signaling axis independent of T-BET (**Fig. 6d**). Nevertheless, the remaining RUNX3 protein levels were insufficient to sustain cytotoxic gene expression in *Tbx21* KO T cells, as GZMB and PRF1 expression were severely reduced **(Fig. 6e)**. Together, these results identify T-BET as a critical mediator of IL-27-induced cytotoxicity, positioning it upstream of RUNX3 in this signaling axis. Moreover, the uncoupling of RUNX3 and T-BET expression in the acute deletion model **(Fig. 6a)** suggests that while RUNX3 is maintained, it alone cannot compensate for the loss of T-BET-driven GZMB production.

**Fig. 6.**
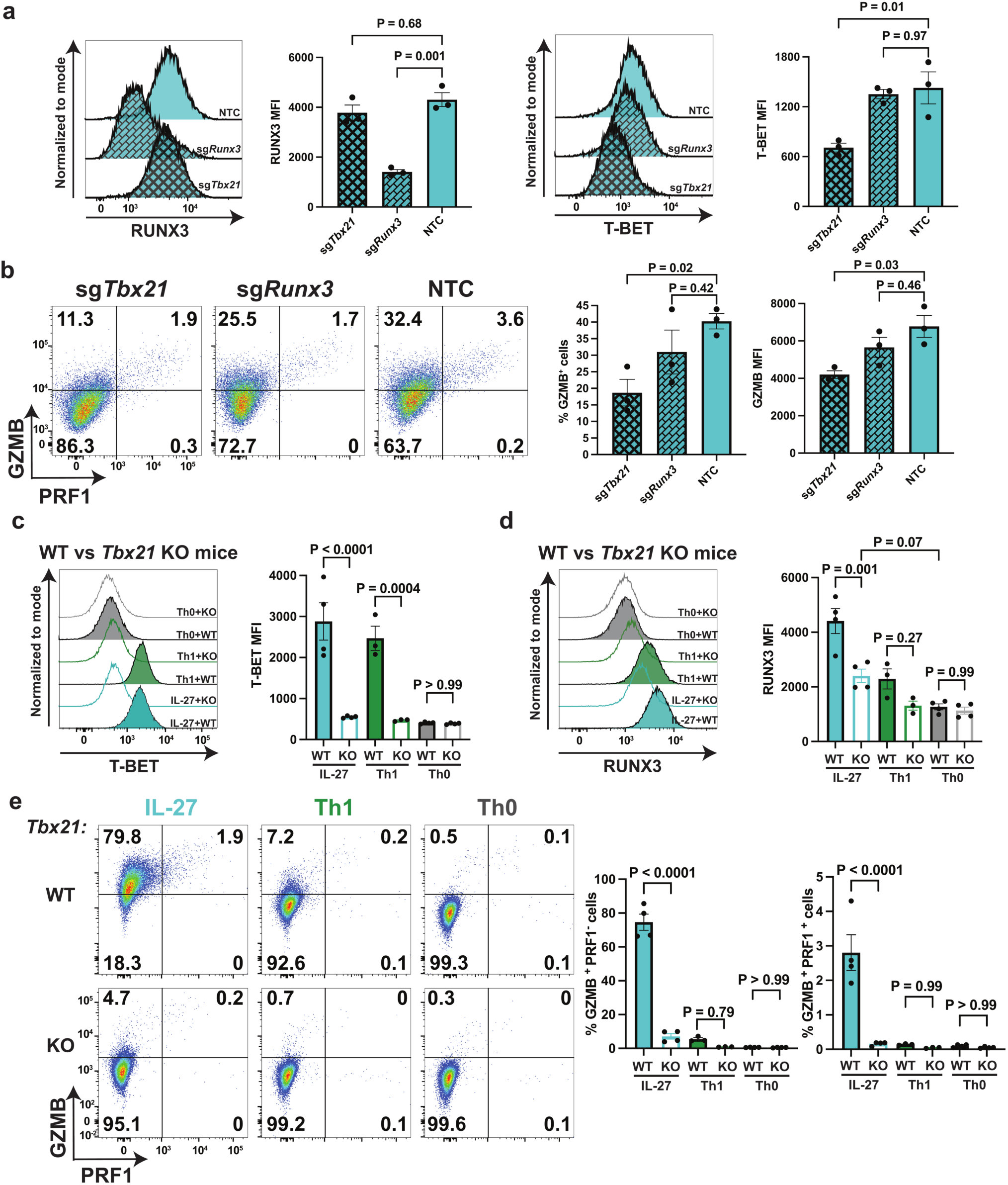
T-BET is essential for the establishment of the IL-27-driven cytotoxic program. **a,b,** *In vitro* genetic deletion of core transcription factors. Naive CD4^+^ T cells from Cas9 mice were activated with α-CD3 and α-CD28 antibodies for 24 hours and transduced with retroviral vectors targeting *Tbx21* (sg*Tbx21*) or *Runx3* (sg*Runx3*), or with a non-targeting control (NTC). 24 hours post-transduction, cells were polarized *in vitro* under IL-27 conditions for 3 days. **a,** Representative flow cytometry histograms and summary bar graphs quantifying the mean fluorescence intensity (MFI) of RUNX3 and T-BET. **b,** Representative flow cytometry plots (left) and summary bar graphs (right) detailing the frequency of GZMB^+^ cells and GZMB MFI across the edited populations. **c–e,** Flow cytometric evaluation of cytotoxic differentiation in wild-type (WT) versus *Tbx21* knockout (KO) CD4^+^ T cells. Naive CD4^+^ T cells from C57BL/6J WT or *Tbx21* KO mice were activated with α-CD3 and α-CD28 antibodies and polarized *in vitro* under the indicated conditions (IL-27, Th1, and Th0) for 3 days. **c,d,** Representative flow cytometry histograms and summary bar graphs quantifying the MFI of T-BET **(c)** and RUNX3 **(d)** across genotypes and polarization conditions. **e,** Representative flow cytometry plots (left) and summary bar graphs (right) detailing the frequencies of GZMB^+^ PRF1^−^ and GZMB^+^ PRF1^+^ populations. Data are presented as mean ± s.e.m. from n = 3–5 independent biological replicates. P values were determined by an ordinary one-way ANOVA with Tukey’s multiple comparisons test.

### IL-27 signaling promotes cytotoxic CD4^+^ T cell differentiation *in vivo* during MCMV infection

CD4-CTLs emerge in non-lymphoid tissues during the acute phase of murine cytomegalovirus (MCMV) infection, which expands in the liver, lungs, and salivary glands^42^. Data from the Human Protein Atlas identifies the liver as the primary anatomical site of baseline *IL-27* mRNA expression **(Extended Data Fig. 7a)**. Given this expression pattern, we asked whether IL-27 signaling is required for CD4-CTL differentiation in the liver during the initial expansion phase of MCMV. We crossed *Il27ra^fl/fl^* mice with *Cd4-Cre* mice to generate a conditional knockout strain (**Fig. 7a**). Conventional CD4^+^ T cells, Treg cells, and CD8^+^ T cells were identified by flow cytometry **(Extended Data Fig. 7b)**. Frequencies and normalized cell counts for CD4^+^ and CD8^+^ T cell populations remained comparable between genotypes, suggesting no broad changes in T cell expansion or survival (**Extended Data Fig. 7c–d**).

**Fig. 7.**
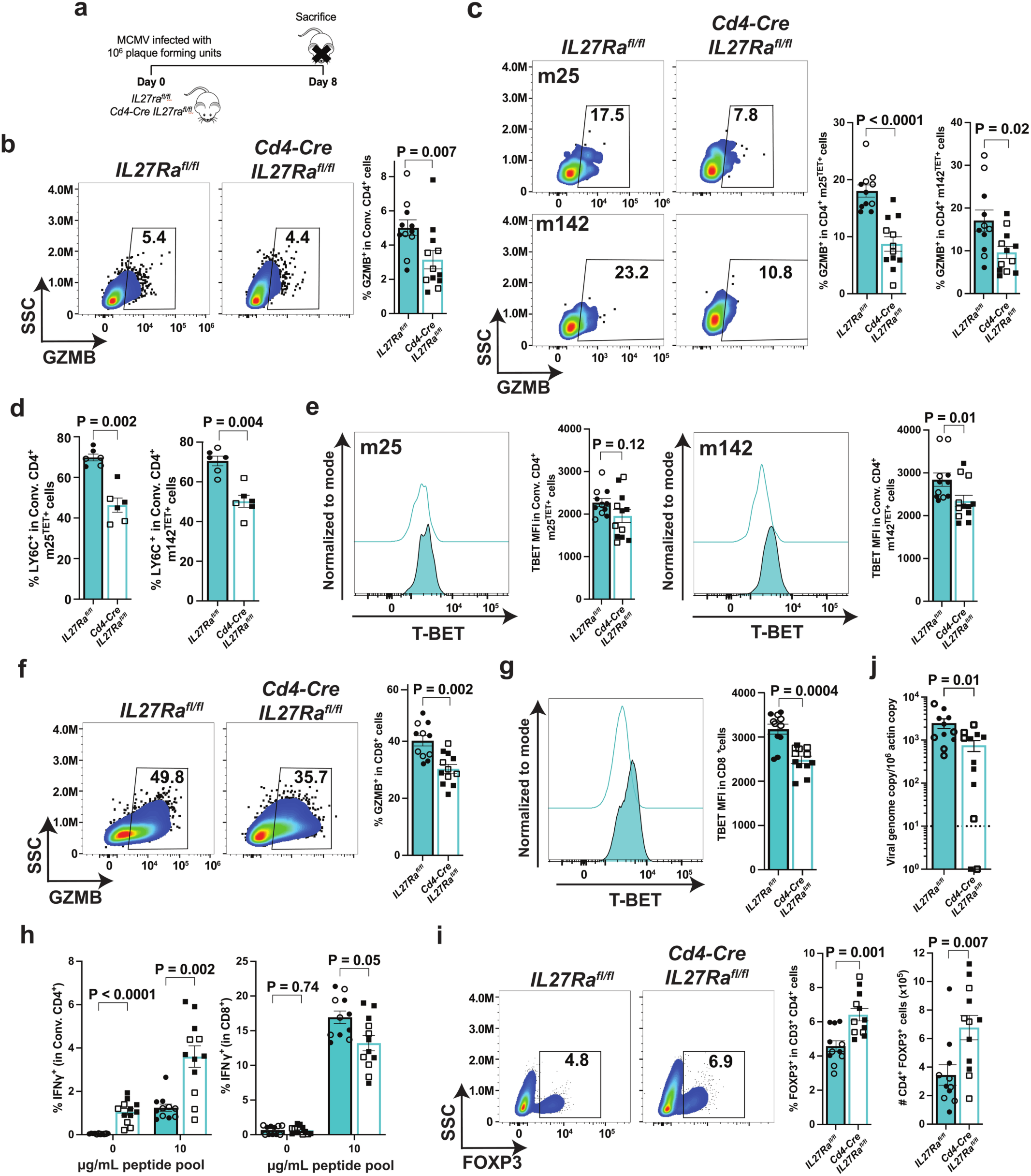
IL-27 receptor signaling *in vivo* is required for cytotoxic CD4^+^ T cell differentiation during MCMV infection. **a,** Schematic of the experimental design. *Il27ra*^fl/fl^ (control) and *Cd4-Cre Il27ra*^fl/fl^ (conditional knockout) mice were infected with 10^6^ plaque-forming units (pfu) of the Smith strain of murine cytomegalovirus (MCMV) and analyzed at day 8 post-infection. Panels **b–j** show cells isolated from the liver. **b,** Representative flow cytometry plots (left) and summary bar graph (right) of intracellular GZMB^+^ population frequencies in conventional CD4^+^ T cells. **c,** Representative flow cytometry plots and summary bar graphs detailing the frequency of GZMB^+^ cells among MCMV-specific m25^+^ (top) and m142^+^ (bottom) CD4^+^ T cells. **d,** Frequencies of LY6C^+^ cells among MCMV-specific m25^+^ (m25^TET+^,left) and m142^+^ (m142^TET+^,right) CD4^+^ T cells. **e,** Representative histograms and summary bar graphs quantifying T-BET mean fluorescence intensity (MFI) in m25-specific CD4^+^ T cells (m25^TET+^,left) and m142-specific CD4^+^ T cells ((m142^TET+^,right). **f,** Flow cytometric evaluation of GZMB^+^ frequency in total CD8^+^ T cells. **g,** Flow cytometric evaluation of T-BET levels in total CD8^+^ T cells. **h,** Summary bar graphs assessing the frequency of IFNγ^+^ cells in conventional CD4^+^ (left) and CD8^+^ (right) T cells following *ex vivo* restimulation with 0 or 10 µg/mL of an MCMV peptide pool (m25, m45, m142, m09, IE3, m38). **i,** Representative flow cytometry plots and summary bar graphs detailing the frequency (left) and absolute number (right) of CD4^+^ FOXP3^+^ regulatory T cells. **j,** Quantification of viral glycoprotein B (gB) DNA copies normalized to actin in the liver from *Il27ra^f^*^l/fl^ and *Cd4-Cre Il27ra*^fl/fl^ mice. Dashed line indicates lower limit of quantitation. Data are presented as mean ± s.e.m. from n = 11 (*Il27ra*^fl/fl^) and n = 12 (*Cd4-Cre Il27ra*^fl/fl^) independent mouse replicates, except for panel **d**, where n = 6 per genotype. Solid symbols denote female mice and open symbols denote male mice. P values were determined by a two-tailed Mann-Whitney test.

In the total CD4 compartment, the loss of IL-27 signaling resulted in a small but significant decrease in the frequency of GZMB^+^ cells, consistent with CD4-CTLs representing a small proportion of total CD4s^5^ (**Fig. 7b**). To investigate the antigen-specific response, we analyzed the immunodominant m25 tetramer^+^(m25^TET+^) and m142 tetramer^+^ (m142^TET+^) MCMV-specific CD4^+^ T cell clones, whose percentages and cell numbers did not change between genotypes (**Extended Data Fig. 7e**). Within both of these antigen-specific T cell pools, we observed a significant reduction in GZMB^+^ frequencies specifically within the CD4^+^ liver populations (**Fig. 7c**), but not in the spleen (**Extended Data Fig. 7f**). The frequency of antigen-specific CD4^+^ T cells expressing LY6C, a putative IL-27-specific surface marker identified in our proteomic comparisons (**Fig. 1f and 3c**), also significantly decreased in *Il27ra* KO mice (**Fig. 7d**). No drop of LY6C was seen in the conventional CD4^+^ T cell pool or any CD8^+^ populations (**Extended Data Fig. 7g**). T-BET protein levels remained stable in conventional CD4^+^ T cells (**Extended Data Fig. 7h**), whereas the m142, but not m25 antigen-specific CD4^+^ T cells, showed decreased TBET levels in the absence of IL-27 receptor signaling (**Fig. 7e**). These data suggest that IL-27 signaling contributes to the induction of liver resident LY6C^+^ and GZMB^+^ CD4-CTLs during MCMV infection.

While Cre expression is driven by the *Cd4* promoter, IL27RA loss in this model occurs at the double-positive thymic stage, deleting the receptor in both CD4^+^ and CD8^+^ T cell populations. The percentages and frequencies of CD8^+^ m45 tetramer^+^ (m45^TET+^) and CD8^+^ m38 tetramer^+^ (m38^TET+^) cells remained the same between genotypes, except for a significant drop in the percentage of CD8^+^ m38^TET+^ cells in KO mice **(Extended Data Fig. 7i)**. The frequency of GZMB^+^ cells within the total CD8^+^ T cell pool was significantly reduced **(Fig. 7f)** consistent with previous cytolytic results in CD8-mediated cancer immunotherapy^20,43^. Though GZMB^+^ frequencies among the m38^TET+^ and m45^TET+^ CD8^+^ T cell clones were unaffected **(Extended Data Fig. 7j).** Similar to our observations in MCMV-specific CD4^+^ T cells, T-BET levels in CD8^+^ T cells were significantly altered by the loss of IL-27 signaling (**Fig. 7g and Extended Data Fig. 7k**).

To evaluate antigen-specific effector capacity, we assessed hepatic CD4^+^ T cells following *ex vivo* restimulation with cognate MCMV peptides. While *IL27ra* KO CD8^+^ T cells showed a subtle but significant defect in IFNγ production, CD4^+^ T cells showed the opposite with a nearly 3-fold increase in IFNγ^+^ cell frequency **(Fig. 7h)** alongside a higher frequency of FOXP3^+^ Treg cells in the liver **(Fig. 7i)**. We next assessed the impact of IL-27 signaling on viral burden in the liver. The reduction of cytotoxic CD4-CTLs via conditional *Cd4*-Cre *Il27ra* knockout resulted in improved viral clearance as assayed by qPCR of MCMV glycoprotein B (gB) DNA copies (**Fig. 7j**), suggesting these cells inhibit anti-viral immunity. Collectively, these findings demonstrate that IL-27 signaling promotes CD4-CTL differentiation *in vivo*, and diverts from Treg and IFNγ^+^ CD4 differentiation during the acute phase of viral infection.

## Discussion

In this study, we identify a unique pathway for CD4-CTL induction, demonstrating that IL-27 acts as a potent, standalone driver of CD4^+^ T cell cytotoxicity. By integrating *in vitro* differentiation, quantitative proteomics, genetic perturbations, and an *in vivo* viral infection model, we establish that IL-27-polarized CD4^+^ T cells settle into a cytotoxic state distinct from the canonical Th1 subset. Both programs depend on the STAT1-T-BET axis, yet they diverge in how this axis is sustained, relying on different signaling and transcriptional mechanisms to maintain their respective identities.

Cytotoxic CD4^+^ T cells arise across diverse immunological contexts^2^, yet the shared expression of subset-defining markers, particularly T-BET and IFNγ has likely obscured their identity. This overlap in lineage marker expression has led investigators to generalize CD4-CTLs as a terminal extension of the Th1 subset^2,6,8,9,12^, masking the possibility of parallel, independently regulated cytotoxic programs. Our mouse and human data begin to disentangle this view. Despite sharing T-BET and IFNγ, IL-27-polarized CD4^+^ T cells and Th1 cells occupy proteomically distinguishable states **(Fig. 2, 3)**. By flow cytometry, GZMB and RUNX3 expression were broadly comparable between human Th1 and IL-27-polarized cells, although IL-27 conditions trended toward a higher frequency of GZMB^+^ PRF1^+^ double-positive cells and higher T-BET abundance **(Extended Data Fig. 2a-d).** IL-27 conditions also showed significantly lower IFNγ, phenocopying the murine data. Nonetheless, the fundamental finding remains that IL-27 drives GZMB induction in both human and murine CD4^+^ T cells.

This IL-27-driven cytotoxic program invites re-evaluation of IL-27’s role in the CD4^+^ T cell compartment, which the field has predominantly characterized as immunoregulatory through induction of IL-10-producing Tr1 cells ^23–25^. This characterization likely reflects, in part, the experimental conditions of earlier studies, which frequently relied on heterogeneous CD4^+^ T cell pools, complex co-culture systems, and variable windows of cytokine exposure^24,25,41,45^. Pairing IL-27 with other environmental signals shapes its outcome. IL-27 alone drives high GZMB and PRF1 expression, while the addition of TGFβ1 represses the acquisition of these cytotoxic features **(Fig. 1b, c)**. Reliance on IL-10 reporter models^46^, where GFP can accumulate, may have further emphasized the link between IL-27 and IL-10 relative to its other functions. Zhang *et al*.^34^ indicate that while IL-27 initiates *Il10* transcription, intracellular IL-10 protein levels are often comparable to those in canonical Th1 subsets. In our own data, the frequency of active IL-10-producing cells in IL-27-driven cultures is low **(Fig. 2e)** but is consistent with previous results^34^. Together, these observations suggest that IL-27’s regulatory and effector functions are not mutually exclusive, and that we have likely underappreciated its capacity to induce cytotoxicity alongside its established role in IL-10 induction.

Our quantitative proteomic analyses defined important differences between the IL-27- and IL-12-driven cell states. IL-27-polarized cells share extensive proteomic similarity with Th1 cells, even enriching more strongly for the established ‘Th1 Cytotoxic Module’ gene set than classical Th1 cells themselves. Yet layered onto this broader overlap is a distinct molecular landscape enriched for cytotoxic machinery and lysosomal degranulation components **(Fig. 2, 3)**. This extensive overlap likely explains why this IL-27-driven population has previously been conflated with canonical Th1 cells when only a subset of markers are considered. The IL-27-driven cytotoxic program operates independently of the autocrine IFNγ feedback loop that sustains canonical Th1 differentiation ^29,30^ **(Fig. 4)**. Blockade of IFNγ signaling significantly reduced T-BET and RUNX3 expression in Th1 cells, yet left the IL-27-driven cytotoxic program fully intact, indicating that IL-27-polarized and Th1 cells rely on distinct regulatory circuits rather than a single program driven by convergent signals. By bypassing the IFNγ requirement, IL-27 establishes a separate differentiation axis for CD4-CTL induction.

Our *in vivo* observations during MCMV infection provide physiological validation for this IL-27-driven cytotoxic program **(Fig. 7)**. Using *Cd4*-Cre-mediated deletion of *Il27ra*, we found that loss of IL-27 receptor signaling significantly decreased GZMB expression in total conventional and antigen-specific, hepatic CD4^+^ T cells (**Fig. 7b-c**). The low magnitude of CD4^+^ GZMB^+^ loss is consistent with CD4-CTLs representing a minority subset of the broader CD4^+^ compartment^5^. Without IL27RA on CD4^+^ T cells, the hepatic compartment showed increased polarization toward FOXP3^+^ Tregs **(Fig. 7i)** and a significantly higher frequency of CD4^+^ IFNγ^+^ cells **(Fig. 7h)**. This increase of Tregs and IFNγ^+^ CD4^+^ T cells in IL27RA-deficient CD4^+^ T cells was also accompanied by a reduced frequency of LY6C^+^ CD4^+^ T cells **(Fig. 7d)**, a surface marker we determined by mass spectrometry to be higher on IL-27-polarized CD4^+^ T cells compared to CD4-IETs or Th1s (**Fig. 1f and 3c**). Although LY6C has been previously suggested as a terminal effector Th1 marker^47^, our data indicate it is not Th1-restricted (**Fig. 1f and 3c**), corroborating IL-27’s established role as an inducer of LY6C expression both *in vitro* and *in vivo*^32^. The loss of GZMB^+^ CD4^+^ T cell in *Cd4*-Cre *Il27ra*^fl/fl^ mice is also consistent with previous work showing that IL-27 receptor signaling expands IL-10^+^ CD4^+^ T cell pool^28^.The resulting hepatic viral reduction in IL27-sufficient mice (**Fig. 7j**) may reflect redundancy in acute antiviral defense, though the long-term impact of losing this dedicated cytotoxic compartment may diverge. As MCMV infection transitions from an acute visceral infection into its characteristic persistent replication phase in the salivary glands, a distinct, late-rising CD4^+^ T cell subset is specifically required to resolve persistent MCMV replication in the salivary gland^44^. Future studies will test whether the IL-27-driven cytotoxic program we describe here contributes to the establishment of this late-rising, protective subset, and whether CD4-CTL loss compromises chronic viral containment in the salivary glands.

At the transcriptional level, both the IL-27-driven and IL-12-driven pathways depend on a shared STAT1-TBET axis **(Fig. 5, 6)**. In the IL-27-driven program RUNX3 plays a secondary role. Although STAT1 partially sustains RUNX3 expression independently of T-BET, CRISPR-mediated deletion of *Runx3* in IL-27-polarized cells indicates that RUNX3 alone is insufficient to drive GZMB induction. Instead, the cytolytic program depends on T-BET. This suggests that RUNX3 is a context-dependent, rather than universal, regulator of cytotoxicity, in contrast to its canonical role as the primary driver in CD8^+^ T cells^13^ and CD4-IETs^3,4^. This divergence extends to a second hallmark of CD4-IET differentiation, which the IL-27-driven program also bypasses. In line with recent reports^31,48^, our data reiterate that ThPOK loss is not a requirement for CD4-CTL differentiation, as IL-27 drives cytotoxic differentiation while maintaining high ThPOK levels **(Fig. 1a)**. This strongly supports our IL-27-driven system as a physiologically relevant platform for generating and studying CD4-CTLs, and adds to a growing body of evidence that mature CD4^+^ T cells retain more phenotypic flexibility in the periphery than the ThPOK-RUNX3 axis alone would predict.

In summary, we have defined IL-27 as a driver of CD4^+^ T cell cytotoxicity, a program that follows a unique molecular path despite sharing the STAT1-T-BET axis with Th1 cells. This work redefines how CD4^+^ T cell cytotoxicity is understood and lays the groundwork for a new generation of biomarkers, engineered T cell products, and therapeutic targets in cancer, chronic infection, and autoimmune disease.

## Methods

### Mice

C57BL/6J, B6.129S6-*Tbx21^tm1Glm^*/J (*Tbx21*^−/−^), B6.129S(Cg)-*Stat1^tm1Dlv^*/J (*Stat1*^−/−^) mice were obtained from Jackson Laboratories. *Zbtb7b^GFP^ Runx3^tdTomato^ mice were obtained from the* Cheroutre lab at LJI and *Cd4-Cre* and *Il27rafl/fl mice*^49^ were obtained from the Li-Fang Lu lab at UCSD. Both male and female mice were used for studies. Mice aged 8 to 12 weeks were used for *in vitro* experiments, while mice aged 10 to 25 weeks were used for the *in vivo* MCMV challenge. All mice were bred and/or maintained in the animal facility at the La Jolla Institute for Immunology (LJI). All experiments were performed in compliance with the study protocol approved by the LJI Institutional Animal Care and Use Committee (IACUC) regulations.

### Mouse naïve CD4^+^ and CD8^+^ T cell isolation, activation and differentiation

Mouse lymph nodes and spleen were harvested and naïve CD4⁺ and CD8⁺ T cells were purified using the EasySep mouse naïve CD4^+^ T Cell Isolation kit and the EasySep mouse naïve CD8⁺ T Cell Isolation kit (STEMCELL technologies), respectively, according to manufacture instructions. T cell activation (0.5 x10^6^ mL; 200 uL/well) was performed using high-binding non-treated tissue culture 96-well plates (Corning) using plate bound α-CD3 (clone 145-2C11, 2 μg/mL) and soluble α-CD28 (clone 37.51 1 μg/mL) monoclonal antibodies in RPMI 1640 Medium, GlutaMAX™ Supplement (Life Technologies) supplemented with 10% FBS (Omega Scientific), 1% penicillin/streptomycin (GIBCO-Life technologies) and 50 μM β-mercaptoethanol (Sigma). For IL-27, Th1 and Th0-polarizing conditions, mIL-27 (25 ng/mL) or mIL-12 (25 ng/mL) and α-IL-4 monoclonal antibody (clone 11B11, 5 μg/ml) or rhIL-2 (20 ng/uL, [800 U]), α-IFNγ monoclonal antibody (clone XMG1.2, 5 μg/ml) and α-IL-4 monoclonal antibody (clone 4B11, 5 μg/ml), respectively, were added to the cultures. Naïve CD8⁺ T cells were cultured in the presence of rhIL-2 (20 ng/uL, 800U). When cells were in culture for more than 3 days, cells were split and rhIL-2 (20 ng/uL, 800U) was added in all polarizing conditions). When indicated, α-IFNγ monoclonal antibody (5 μg/mL), α-IL-21 monoclonal antibody (Clone FFA21, 5 μg/mL), hTGFβ (2.5 ng/mL), RA (25 nM) or STATTIC (2 µM) were added. For cultures requiring STATTIC, cells were incubated with the inhibitor for 40 min prior to activation and cytokine addition. Cells were maintained in a standard tissue culture incubator containing 5% CO_2_.

### Human naïve CD4^+^ T cell isolation, *in vitro* activation and differentiation

Peripheral blood mononuclear cells (PBMCs) were isolated from whole blood of healthy donors via Ficoll-Paque Plus (Sigma-Aldrich) density gradient centrifugation. Primary human naive CD4^+^ T cells were subsequently enriched using the EasySep Human Naive CD4^+^ T Cell Isolation Kit II (STEMCELL Technologies) according to the manufacturer’s instructions, with a resulting purity of > 90% routinely confirmed by flow cytometry.

Isolated naive T cells were cultured in RPMI 1640 Medium supplemented with GlutaMAX, 10% FBS (Omega Scientific), and 1% penicillin/streptomycin (GIBCO). Cells were seeded at a density of 0.5 x10^6^ cells per mL (1mL per well) in 24-well non-treated tissue culture plates (ThermoFisher) and activated using plate-bound α-CD3 (clone OKT3; 1 ug/mL) and α-CD28 (clone 9.3; 1 ug/mL) monoclonal antibodies.

For *in vitro* polarization, cells were cultured under the following conditions: IL-27 conditions (hIL-27, 25 ng/mL); Th1 conditions (rhIL-12, 25 ng/mL and α-IL-4, 5 ug/mL); or Th0 control conditions (hIL-2, 20 ng/mL [800 U], α-IFNγ, 5 ug/mL, and α-IL-4, 5 ug/mL). All human blood sample protocols were approved by the La Jolla Institute for Immunology Institutional Review Board and the Normal Blood Donor Program.

### Flow cytometry

A range of 2e5 – 1.5e6 cells were used to perform flow cytometry phenotype assessment. Expression of surface markers was evaluated using the appropriate fluorochrome-conjugated monoclonal antibodies and viability addressed using fixable viability dye in a total volume of 50 μL. Cells were incubated in the dark for 20 minutes in PBS containing 2% FBS at 4°C and then washed once in the same medium at 500 g for 5 minutes. For cytokine staining, 4e5 – 1.5e6 cells were previously stimulated with phorbol 12-myristate 13-acetate (PMA) (100 ng/ml, Sigma) and ionomycin (1 μg/ml, Sigma) in the presence of GolgiStop (BD Biosciences), as indicated by the manufacture in complete RPMI media for 3.5h at 37°C (total volume of 1 mL). For intracellular staining of transcription factors and cytokines fixation and permeabilization was done using the Foxp3/Transcription Factor intracellular staining kit from (BD Biosciences) as per the manufacturer’s instructions and cells were incubated for 40 minutes at 4°C. In experiments using *Zbtb7b^GFP^ Runx3^tdTomato^* mice, cells were previously fixed with BD cytofix for 20 min to maintain GFP signal. Cells were then washed with the permeabilization buffer and incubated with intracellular antibodies for 1 hour at RT followed by 4°C overnight incubation. Cell analyses were performed on Fortessa cytometer (BD Biosciences) and Aurora (Cytek). A minimum of 20,000-30,000 events (viability dye– negative) were recorded for each sample. Data analyses were performed using FlowJo software (Tree Star, Ashland, OR).

### Antibodies

Antibodies used for flow cytometry are listed below, grouped by experimental context.

#### *In vitro* Mouse T cell experiments

Ghost Dye (viability)-UV450, CD4-BV711 (GK1.5) Granzyme B-Pacific Blue (GB11), Perforin-APC (S16009A), FOXP3-PE-Cy7 (FJK-16s), T-BET-PerCP-Cyanine5.5 (4B10), T-BET-BV785 (4B10), RUNX3-PE (R3-5G4), IL-10-BV605 (JES5-16E3), CD8b-BV510 (H35-17.2), Thy1.1-PE-Cy7 (OX-7), IFNγ-BUV737 (XMG1.2), pSTAT1 (pY701)-BV421 (4a), pSTAT3 (pY705)-AF488 (4/P-STAT3), pSTAT4 (pY693)-PerCP-Cyanine5.5 (38/p-Stat4), and pSTAT5 (pY694)-AF647 (47/Stat5(pY694)).

#### *In vitro* Human T cell experiments

CD4-FITC (RPA-T4), CD25-BUV395 (M-A251), CD45RA-PerCP-Cyanine5.5 (HI100), CD62L-BV510 (DREG-56), T-BET-BV785 (4B10), RUNX3-PE (R3-5G4), IFNγ-PE-Cyanine7 (B27), Granzyme B-Pacific Blue (GB11), Perforin-APC (dG9), and Ghost Dye (viability)-UV450.

### Phospho-STATs induction and PhosFlow staining

Naïve CD4^+^ T cells were incubated in cold PBS containing 2% FBS with α-CD3 (clone 145-2C11, 1 μg/mL) and α-CD28 (clone PV1, 1 μg/mL) per million of naïve CD4⁺ T cells for 20 minutes at 4°C. Armenian hamster crosslinker polyclonal antibodies (Jackson ImmunoResearch) were then added at a final concentration of 18 μg/mL and incubated during 20 minutes at 4°C. After incubations, cold PBS was added and cells were collected by centrifugation. Naïve CD4⁺ T cells were resuspended to a concentration of 1e6 cells/mL in 37°C PBS containing 2% FBS with either mIL-27 (25 ng/mL) or mIL-12 (25 ng/mL) or no cytokines as a control (200 uL PBS-FBS buffer). After 30 minutes, 200 μL of 37°C BD Cytofix buffer (BD biosciences) were directly added and cells incubated during 10 minutes at 37°C. After centrifugation, 1 mL of BD Phosflow Perm Buffer III (BD biosciences) was added and cells incubated during 30 minutes at 4°C. After incubation, cells were washed twice with cold PBS containing 2% FBS and pSTAT fluorochrome-conjugated monoclonal antibodies added and incubated during 45 minutes at RT. Cells were washed with cold PBS containing 2% FBS before analysis in the flow cytometer.

### TMTpro mass spectrometry-based quantitative proteomics

Naive CD4^+^ T cells were isolated from *Zbtb7b^GFP^ Runx3^tdTomato^* reporter mice and differentiated as described above. Following confirmation of subset differentiation by flow cytometry, previously snap frozen pellet were lysed in a buffer containing 8 M urea, 50 mM Tris (pH 8.0), 75 mM NaCl, 1 mM EDTA, and a 1X protease and phosphatase inhibitor cocktail. Protein concentration was quantified via Bradford assay. Proteins were reduced with 10 mM TCEP, alkylated with 5 mM iodoacetamide, and digested sequentially with Lys-C (Fuji) for 2 h followed by overnight digestion with proteomics-grade trypsin (Promega). Digested peptides were acidified to 1% formic acid and subjected to on-column TMT-labeling^50^. Briefly, peptides were trapped on C18 resin (EMPORE) and labeled directly on the column using TMTpro 18-plex reagents (channels 126 to 135) as described in^50^ with the exception that 50 mM phosphate buffer pH 8.0 was used instead of HEPES. The multiplexed TMTpro sample was pooled and fractionated via StageTip fractionation into 11 fractions using a stepwise acetonitrile gradient in 20 mM ammonium formate, with the first two fractions concatenated to yield 10 fractions for analysis. Fractions were analyzed by liquid chromatography-tandem mass spectrometry (LC-MS/MS) using ultra-high-pressure liquid chromatography (UHPLC) system coupled online to an Orbitrap Eclipse Tribrid mass spectrometer (Thermo Fisher Scientific). Peptides were separated on an analytical C18 capillary column. Mass spectrometry data acquisition was performed in data-dependent acquisition (DDA) mode. Selected precursor ions were fragmented via higher-energy collisional dissociation (HCD) prior to Orbitrap MS2 mass analysis. Mass spectra were searched and processed using the Spectrum Mill software platform (Agilent and Broad Institute) against the UniProt mouse database (12/28/2017). Reporter ion intensities were corrected for isotopic impurities using manufacturer-supplied lot-specific correction factors within Spectrum Mill. Downstream data normalization, statistical differential expression analysis, and quality control were performed using Protigy (https://github.com/broadinstitute/protigy), and final relative protein abundance matrices were visualized using Morpheus (https://software.broadinstitute.org/morpheus/).

### Hierarchical clustering, K clustering, Marker selection and protein correlation analysis

Hierarchical clustering, k-means clustering, marker selection, and protein correlation analyses were performed using the Morpheus web platform (Broad Institute; https://software.broadinstitute.org/morpheus/). Both hierarchical, k-means clustering and protein correlation analyses were restricted strictly to differentially abundant proteins, with significance for differential abundance determined based on F-statistics (adjusted P value ≤ 0.050). Hierarchical clustering was applied to rows and/or columns as indicated by the respective dendrograms for individual analyses, utilizing one minus Pearson’s correlation as the distance metric. For marker selection, a signal-to-noise metric was utilized in an iterative one-versus-all approach. Significance was determined utilizing 1,000 permutations, and only proteins with a nominal P ≤ 0.01 were retained for downstream analysis. For protein correlation analysis, one minus Pearson’s correlation was utilized as the distance metric. Differentially abundant proteins demonstrating a Pearson correlation coefficient of ≥ 0.75 with TBX21, RUNX3, and GZMB were selected. The final reported protein set was defined by the overlap (intersection) of the proteins highly correlated with all three of these proteins.

### Pathway and Process enrichment analysis

Pathway and process enrichment analysis was performed using the Metascape web portal (https://metascape.org)^51^. The lists of selected proteins were inputted into the platform, with the input species set to *Mus musculus* and the analysis species set to *Homo sapiens*. Enrichment analysis was conducted utilizing multiple ontology sources, including Gene Ontology (GO) Biological Processes, KEGG Pathways, and Reactome Gene Sets. For statistical significance, terms were required to meet a threshold of a P value < 0.01 alongside default Metascape filtering and clustering parameters. Finally, representative terms for visualization and discussion were manually curated from the resulting statistically significant clusters based on their specific biological relevance to the experimental context.

### Gene Set Enrichment Analysis (GSEA) analysis

Gene Set Enrichment Analysis was performed using the GSEA Preranked tool (Broad Institute). Proteins were pre-ranked based on their signed logP value from the statistical comparison between experimental conditions. The pre-ranked lists were queried against the Molecular Signatures Database (MSigDB, version 2026.1). For the analysis of human gene sets, the C2 (curated gene sets) and C7 (immunologic signature gene sets) collections were utilized. Mouse gene symbols were mapped to human orthologs using the MSigDB remapping chip file (Mouse_Gene_Symbol_Remapping_Human_Orthologs_MSigDB.v2026.1.Hs.chip). For the analysis of mouse gene sets, the M7 collection (immunologic signature gene sets) was utilized alongside the corresponding mouse gene symbol remapping chip (Mouse_Gene_Symbol_Remapping_MSigDB.v2026.1.Mm.chip). Across all analyses, default parameters were used where applicable, including 1,000 permutations and a weighted scoring scheme. Identifiers were collapsed to gene symbols, and the analysis was restricted to gene sets containing a minimum of 15 and a maximum of 500 genes.

### Principal component analysis

Principal component analysis (PCA) was performed in R v4.6.0 (prcomp, centered and scaled) on log2 protein abundances across all proteins quantified in every sample. To analyze gene sets contributing to the resultant projections, the loadings from PC1 and PC2 were analyzed using Gene Set Enrichment Analysis (fgsea v1.38.0, 10,000 permutations, gene set size 10–500). Because MSigDB gene set collections are curated in human, mouse gene symbols were first mapped to human orthologs using the babelgene package v22.9 (HCOP consensus, ≥3 supporting databases, restricted to 1:1 orthologs), and gene sets were retrieved with msigdbr v26.1.0. GSEA was run against the C7 immunologic signature collection, and a manually curated set of immunologically relevant significant signatures (adj. p ≤ 0.05) was displayed with their respective normalized enrichment score (NES), positioned according to the directionality of the sample clusters along PC1 and PC2.

### Human Protein Expression Analysis

To evaluate the baseline tissue distribution of human IL-27, publicly available expression data was retrieved from the Human Protein Atlas (HPA; https://www.proteinatlas.org/). The target "IL-27" was queried, and protein expression profiles across normal human tissues were examined utilizing the Tissue Atlas module.

### Cloning of retrovirus sgRNA constructs

Protospacer sequences targeting *Runx3* and *Tbx21* were obtained from the mouse Brie library^52^ using four protospacers per gene, and four non-targeting control protospacers were obtained from^53^ **(Supplementary Table 7)**. For each protospacer, a forward oligonucleotide (5′-CACCG followed by the 20-nt protospacer) and a reverse oligonucleotide (5′-AAAC followed by the reverse complement of the protospacer and a terminal C) were synthesized (IDT, 25 nmol scale), so that the appended 5′ G provides the U6 transcription start nucleotide and the resulting CACC and AAAC overhangs are compatible with BbsI-linearized pSIRG-NGFR^54^. Oligonucleotides were resuspended to 200 µM in nuclease-free water. Forward and reverse oligonucleotides for each protospacer were annealed and phosphorylated in a 10 µL reaction containing 1 µL of each oligonucleotide (200 µM), 1 µL of 10x T4 DNA ligase buffer and 0.5 µL of T4 polynucleotide kinase (NEB) by incubation at 37 °C for 30 min, denaturation at 95 °C for 5 min and ramping to 25 °C at 6 °C min⁻¹, and the annealed oligonucleotides were diluted 1:200 in water. pSIRG-NGFR (5 µg) was digested with BbsI-HF (NEB, 60 U) in 1x rCutSmart buffer (50 µL, 37 °C, 60 min), resolved on a 1% agarose gel, and the 8,485-bp linearized backbone was purified (Monarch Spin DNA Gel Extraction Kit, NEB). Each protospacer was ligated independently into the linearized backbone in a 10 µL reaction containing 25 ng of backbone, 1 µL of the diluted annealed oligonucleotides, 1 µL of 10x T4 DNA ligase buffer and 0.5 µL of T4 DNA ligase (NEB, 400 U µL⁻¹), incubated at room temperature for 10 min and heat-inactivated at 65 °C for 10 min, alongside a no-insert negative control. For each gene, equal volumes of the four protospacer ligations were combined (1 µL each, 4 µL total) and transformed into 100 µL of chemically competent NEB Stable E. coli (prepared in house), plated on a 10-cm LB-ampicillin agar plate and grown overnight at 30 °C. Colonies were scraped into 5 mL of warm LB and pooled and 4.5mLs were expanded in 50 mL total of LB-ampicillin overnight before plasmid purification (ZymoPURE II Plasmid Midiprep Kit), yielding a per-gene minipool of four protospacers. To assess per-sgRNA representation, a 387-bp region spanning the protospacer was amplified from each minipool (primers in **Supplementary Table 7**), gel-purified and analyzed by long-read amplicon sequencing (Plasmidsaurus Premium PCR). For knockout experiments, the *Runx3* and *Tbx21* minipools were compared with the non-targeting control minipool, and minipools were regrown from glycerol stocks for additional experimental replicates.

### Retroviral production and supernatant preparation

Lenti-X 293T cells were seeded at 12e6 cells per T75 flask in complete Opti-MEM, supplemented with 5% FBS (Gibco), 1 mM Sodium Pyruvate (Gibco), and 1x MEM NEAA (Gibco). The following day, cells were transfected using the Lipofectamine 3000 kit in serum-free Opti-MEM. The transfection mixture consisted of 13 µg of MSCV plasmid encoding hCD19 CAR Thy1.1 or pSIRG hGNFR encoding different sgRNAs, and 8.66 µg of the pCL-ECO packaging plasmid. Following a 15-minute room temperature incubation, the transfection complex was added to the cells. Cells were incubated for 4 hours at 37°C and 5% CO2, after which the media was replaced with 15 mL of complete Opti-MEM. Viral supernatants were harvested 18 to 24 hours post-media change. The collected supernatant was centrifuged at 500g for 10 minutes to remove cellular debris and subsequently passed through a 0.45 µm filter. To concentrate the virus, Retro-X concentrator (Takara) was added at a 1:3 ratio to the clarified supernatant and incubated overnight at 4°C. The following day, the mixture was centrifuged at 1500g for 45 minutes at 4°C, and the resulting viral pellets were resuspended in 1.5 mL of RPMI/T75 flask.

### T cell transduction

Non-treated 24-well culture plates were coated overnight at 4°C with 20 µg/mL RetroNectin (Takara) in PBS. Concurrently, naive CD4^+^ T cells were seeded at a density of 0.5 × 10⁶ cells/mL (200 µL per well) in high-protein-binding, non-tissue-culture-treated 96-well plates (Corning). These cells were stimulated overnight using plate-bound α-CD3 (clone 145-2C11, 2 µg/mL) and soluble α-CD28 (clone 37.51, 1 µg/mL) antibodies in RPMI 1640 medium supplemented with GlutaMAX™ (Life Technologies), 10% FBS (Omega Scientific), 1% penicillin-streptomycin (Gibco), and 50 µM β-mercaptoethanol (Sigma).

The following day, the RetroNectin-coated plates were blocked with a 2% BSA solution in PBS for 30 minutes at room temperature, followed by a PBS wash. Concentrated viral supernatant (500 µL/well) was immediately added to the washed plates. To facilitate viral binding to the RetroNectin, the plates were subjected to spinoculation at 2000g for 2 hours at 32°C.

After spinoculation, the viral supernatant was removed from the plates, and 500 µL of the activated T cell suspension (0.5 × 10⁶ cells total) was added directly to each well. The cells were spinoculated at 2000g for 1.5 hours at 32°C and then incubated for an additional 3 to 4 hours to maximize infection efficiency. Following transduction, the T cells were collected, washed with PBS, and cultured.

### *In vitro* cytotoxic assay

Naïve CD4^+^ T cells were stimulated overnight with plate-bound α-CD3 antibody (clone 145-2C11, 2 µg/mL) and soluble α-CD28 antibody (clone 37.51, 1 µg/mL) in RPMI 1640 medium supplemented with GlutaMAX™ (Life Technologies), 10% fetal bovine serum (FBS; Omega Scientific), 1% penicillin–streptomycin (Gibco), and 50 µM β-mercaptoethanol (Sigma). Following activation, cells were retroviral transduced as described above. After transduction, cells were then maintained for 3 days under two parallel polarization conditions: IL-27 (mIL-27 - 25 ng/mL) or Th1 (mIL-12-25ng/mL and α-IL-4 monoclonal antibody at 5 μg/ml) in complete RPMI medium prior to co-culture with target cells. Transduction efficiency of ≥80% was confirmed by flow cytometry by assessing THY1.1 surface level expression (a surrogate marker for CAR expression), along with a viability dye to exclude dead cells.

For co-culture assays, activated α-hCD19 CD4^+^ CAR T cells (effectors) were incubated with Raji B cells constitutively expressing luciferase (targets) in complete RPMI at an effector-to-target (E:T) ratio of 10:1. Target Raji cells (>95% viable) were seeded into round-bottom 96-well plates at a density of 5,000 cells per well in 50 uL of complete RPMI medium. Effector CAR T cells were added at a density of 50,000 cells per well in an equal volume, and the co-culture plate was centrifuged at 300 x g for 5 min before being incubated at 37°C with 5% CO2 for 48 h.

Target cell lysis was quantified using the Britelite Plus Reporter Gene Assay System (Revvity) according to the manufacturer’s instructions. Briefly, after equilibrating the co-culture plate to room temperature, 50 uL of the cell suspension was transferred to a 96-well white flat-bottom CulturPlate (Revvity), followed by the addition of 50 uL of reconstituted Britelite Plus luciferase reagent. Plates were agitated at 150 rpm for 1 min, and endpoint luminescence was recorded using a Revvity EnSight multimode plate reader.

### *In vivo* MCMV infection

MCMV (Smith strain) was originally produced in 3T3 cells from cloned and sequenced BAC DNA, then amplified *in vitro* in 3T3 cells and titrated on M2-B24 cells (murine fibroblast-like cells). Male and female mice were infected i.p. with 10^6^ pfu in 100μL of PBS and euthanized 8-days post infection. Following euthanasia by CO_2_ inhalation, mice were perfused with cold PBS through the heart left-ventricle. Upon spleen and liver harvesting, a small piece of tissue was flash-frozen in liquid nitrogen and stored at −80°C for DNA extraction and viral load quantification by qPCR, as previously described^55^. Spleens and livers were then processed by mechanical dissociation, RBC lysis, and density gradient (liver only), as previously detailed^56^. Single-cell suspensions were either directly stained with NIH-provided MHC-tetramer, followed by surface and intracellular staining or restimulated for 4 hours with 10 µg/mL of MCMV-derived antigens (m09_133-147_, m25_409-423_, m142_24-38_, IE3_416-423_, m38_316-323_, m45_985-993_, in presence of brefeldin A, prior to cytometric staining.

Antibodies used for flow cytometry in MCMV experiments were: NK1.1-BV510 (PK136), CD3-BUV615 (17A2), CD4-BUV395 (GK1.5), CD8a-BUV805 (53-6.7), Granzyme B-AF700 (QA16A02), Granzyme B-PE-Cy7 (QA16A02), Perforin-PE/Dazzle594 (S16009A), FOXP3-AF488 (MF-14), T-BET-BV785 (4B10), and IFNγ-eFluor 450 (XMG1.2), together with Fixable Viability Dye-eFluor 506 (Blue) and PE- or APC-conjugated streptavidin. Biotinylated I-Ab-restricted (m09, GYLYIYPSAGNSFDL; m25, NHLYETPISATAMVI; m142, RSRYLTAAAVTAVLQ) and H2-Kb/H2-Db-restricted (m38, SSPPMFRV; m45, HGIRNASFI; IE3, RALEYKNL) peptide-MHC monomers were obtained from the NIH Tetramer Core Facility.

### Statistical analyses

All statistical comparisons were performed in GraphPad Prism V11. Unpaired t-tests or Mann-Whitney tests were used for pairwise comparisons, and ordinary one-way ANOVA with Tukey’s post-hoc correction was used for comparisons involving more than two groups. P values are only shown for relevant comparisons.

## Author contributions

M.I.M. and S.A.M. conceived and designed the study. M.I.M., S.F.B. and M.J. performed *in vitro* mouse T cell polarization experiments. M.I.M. performed proteomic profiling, retroviral production, supernatant preparation, T cell transduction and CRISPR-based mechanistic experiments. L.S.S., K.M., H.L. and G.A. performed molecular cloning. R.M. performed the MCMV *in vivo* experiments under the supervision of C.B. K.M. performed the *in vitro* human CD4^+^ T cell experiments and the *in vitro* cytotoxic assay. M.I.M., L.S.S. and S.A.M performed bioinformatic and proteomic data analysis. E.E. provided the CAR construct. Y.Z. provided the pSIRG construct and invaluable insight. S.A.M. supervised the project. M.I.M and S.A.M. acquired funding for the study. M.I.M. and S.A.M. wrote the manuscript. All authors provided feedback on the manuscript.

## Acknowledgments

We thank Anjana Rao for the CAR construct. We thank Li-Fang Lu for the *Cd4-Cre Il27ra^fl/fl^* mice. We thank Mitch Kronenberg, Michael Croft, Pandurangan Vijayanand, B. Hamilton, Egle Kvedaraite, and Roberta Nowak for useful discussion, and Sandy Boek Werness and Bruce Werness for their support. We thank Peter Jones, Ana Causton, Nicola Eaton, and Erik Leksell of the Department of Laboratory Animal Care at LJI for assistance with mouse colonies. We thank the Flow Cytometry Core Facility at LJI for their assistance. This work was supported by the Global Autoimmune Institute, Curebound Discovery Grant 23DG08 and The Tullie and Rickey Families SPARK Awards for Innovations in Immunology.

## Data availability

The mass spectrometry-based proteomics data generated in this study have been deposited in the MassIVE repository under accession number MSV000102431 and will be made publicly available at ftp://massive-ftp.ucsd.edu/v13/MSV000102431/ upon publication. Supplementary tables, source data underlying the figures, and the mouse lines and reagents generated in this study, are available from the corresponding author upon request.

## Competing interests

The authors declare no competing interests.

---

**Extended Data Fig. 1.**
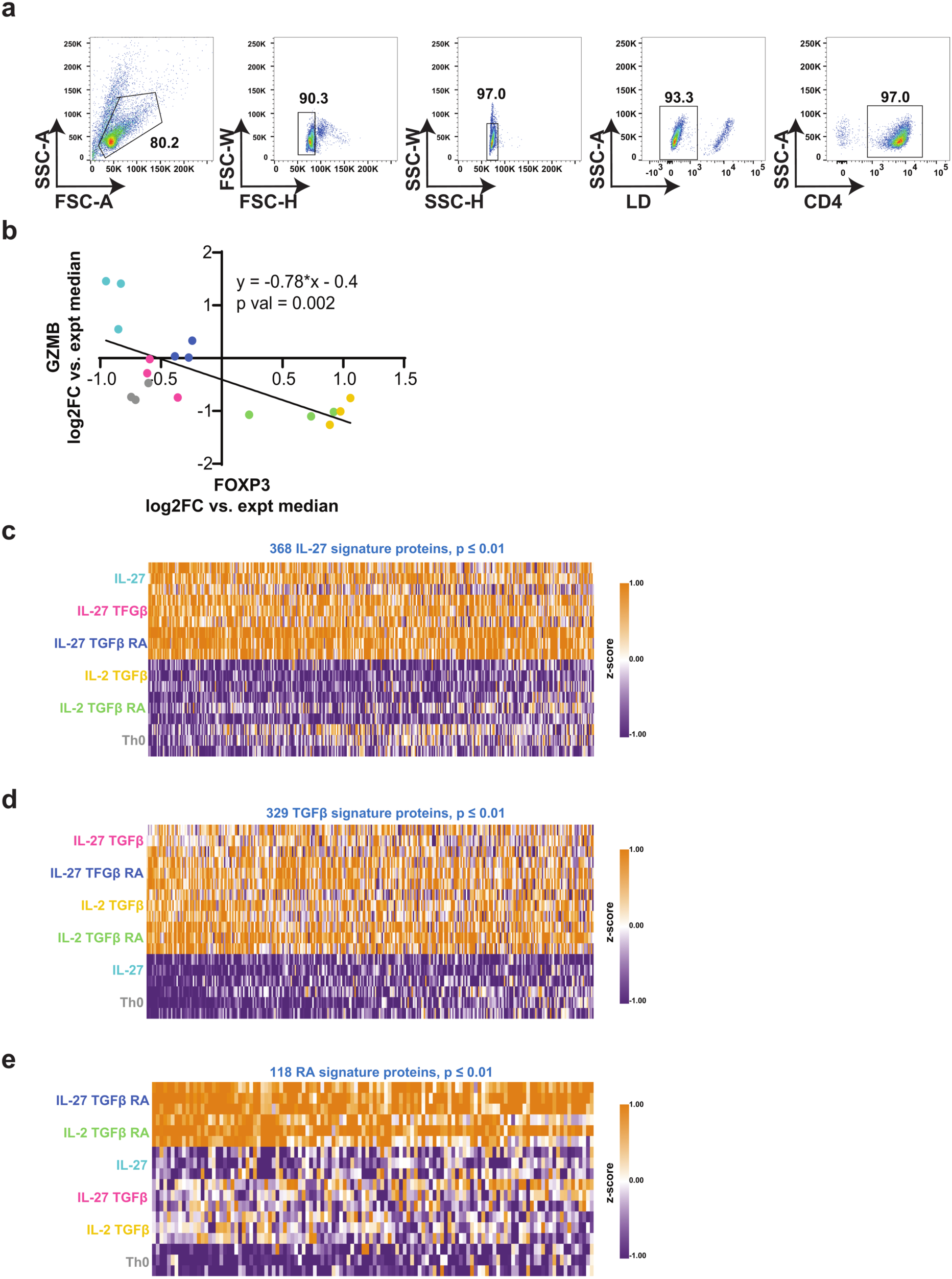
Proteomic profiling and marker selection analysis of polarized CD4^+^ T cells. Naive CD4^+^ T cells from *Zbtb7b*^Gfp^ *Runx3*^tdTomato^ mice were activated with α-CD3 and α-CD28 antibodies and polarized *in vitro* under the indicated conditions for 6 days prior to mass spectrometry-based proteomic analysis. **a,** Gating strategy for the analysis of live CD4^+^ T cells. **b,** Scatter plot detailing the correlation between GZMB and FOXP3 relative protein abundance levels (log2 fold change versus experiment median). The solid line represents the linear regression (y = −0.78x - 0.4, P = 0.002). **c–e,** Marker selection analysis (p ≤ 0.01) identifying distinct proteomic main-effect signatures driven by IL-27 **(c)**, TGFβ **(d)**, or retinoic acid (RA) **(e)**. Data in b–e are derived from n = 3 independent experiments.

**Extended Data Fig. 2.**
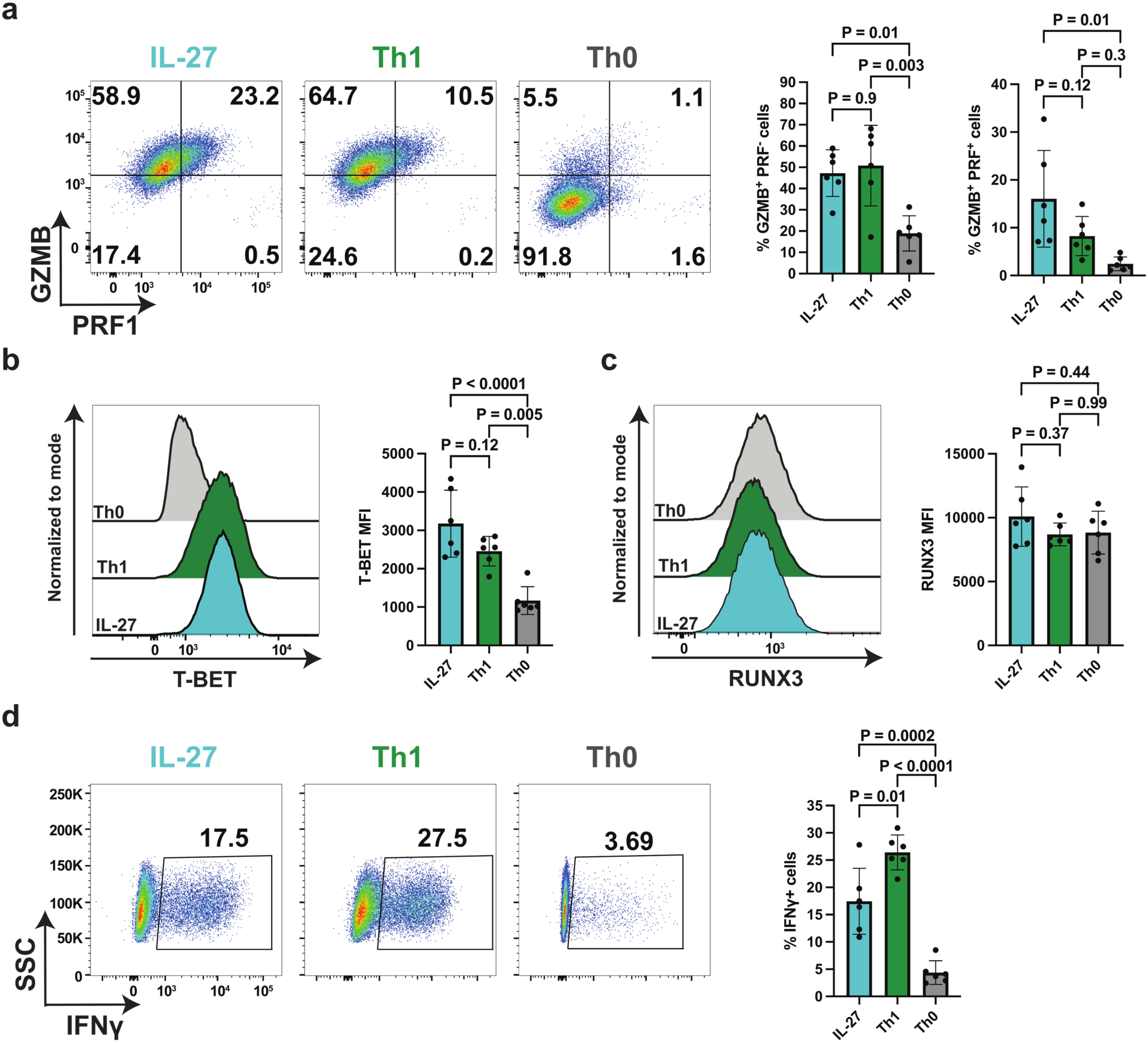
Phenotypic characterization of IL-27-polarized primary human CD4^+^ T cells. Naïve CD4^+^ cells were activated with α-CD3 and α-CD28 antibodies and polarized *in vitro* under IL-27, canonical Th1, or unpolarized (Th0) conditions for 3 days. **a,** Representative flow cytometry plots (left) and summary quantitative bar graphs (right) detailing the frequencies of intracellular Granzyme B (GZMB) and Perforin (PRF1) single-positive (GZMB^+^ PRF1^-^) and double-positive (GZMB^+^ PRF1^+^) populations. **b,c,** Representative flow cytometry histograms and summary bar graphs of the mean fluorescence intensity (MFI) for the transcription factors T-BET **(b)** and RUNX3 **(c)**. **d,** Representative flow cytometry plots (left) and summary quantification (right) of intracellular IFNγ expression. For all panels, data are presented as mean ± s.e.m. from n = 6 independent biological replicates from 5 different donors. P values were determined by ordinary one-way ANOVA with Tukey’s multiple comparisons test.

**Extended Data Fig. 3.**
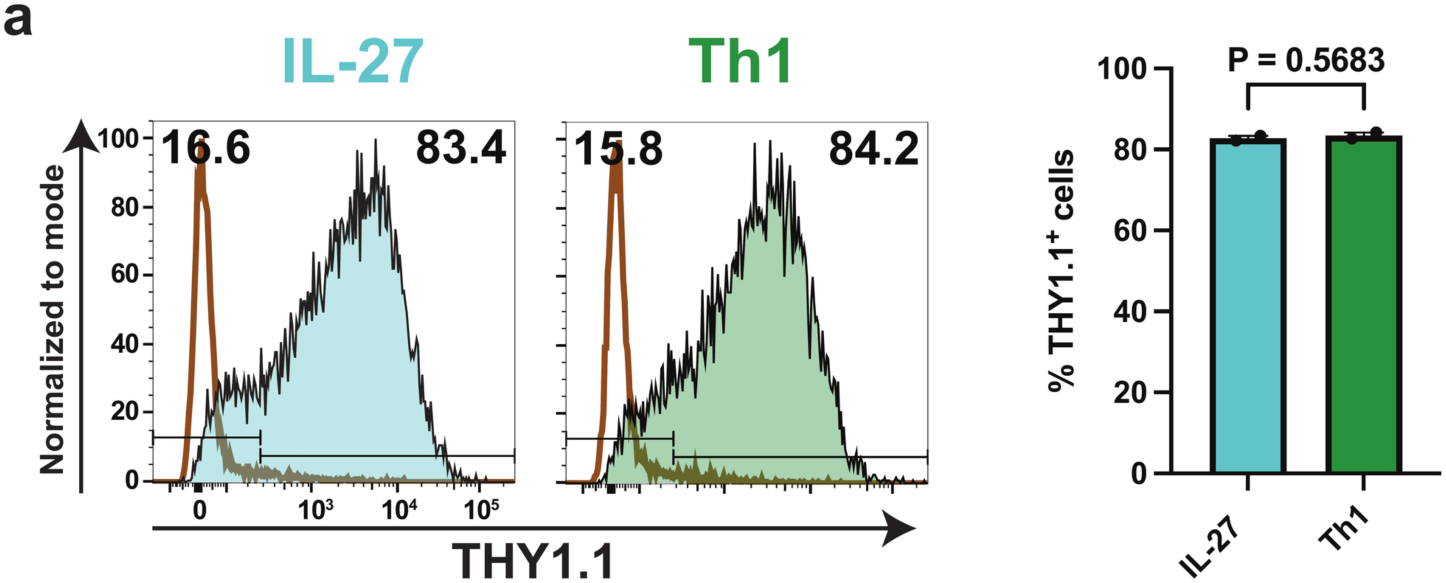
Validation of CAR transduction in primary mouse CD4^+^ T cells. Assessment of chimeric antigen receptor (CAR) transduction efficiency. Naive CD4^+^ T cells from C57BL/6J mice were activated, transduced with a retroviral vector encoding an anti-human CD19 (hCD19) CAR coupled to a THY1.1 reporter, and polarized *in vitro* under IL-27 or Th1 conditions. Representative flow cytometry histograms (left) and summary bar graph (right) detail the transduction efficiency, quantified by the frequency of THY1.1^+^ cells. The brown open histograms represent untransduced control cells. Data are presented as mean ± s.e.m. from n = 2 independent biological experiments. P values were determined by unpaired two-tailed Student’s t-test.

**Extended Data Fig. 4.**
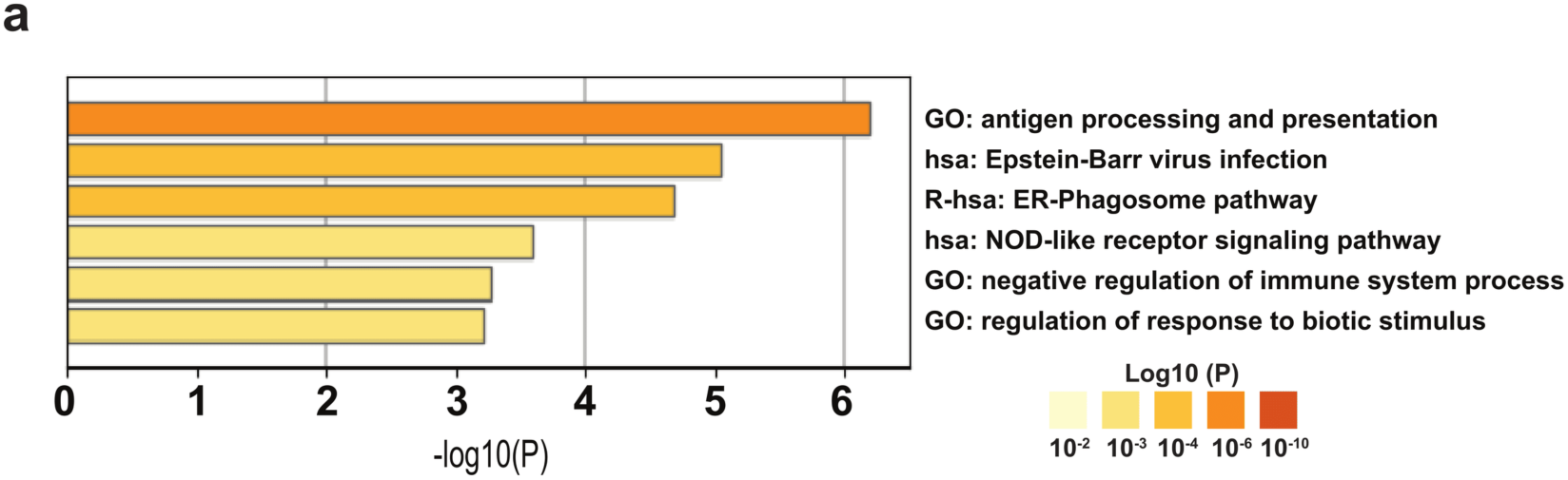
Functional enrichment analysis of the IL-27-driven co-expression module. Bar graph detailing Gene Ontology (GO) and biological pathway enrichment analysis of co-expression module. This module was identified by performing Pearson’s correlation analysis of the differentially abundant proteins (Fig. 3b) to isolate proteins that significantly correlated (r ≥ 0.75) with the expression of GZMB, RUNX3, and TBX21 (as defined in Fig. 3e). The top enriched functional pathways are displayed and ranked by –log10(P) value.

**Extended Data Fig. 5.**
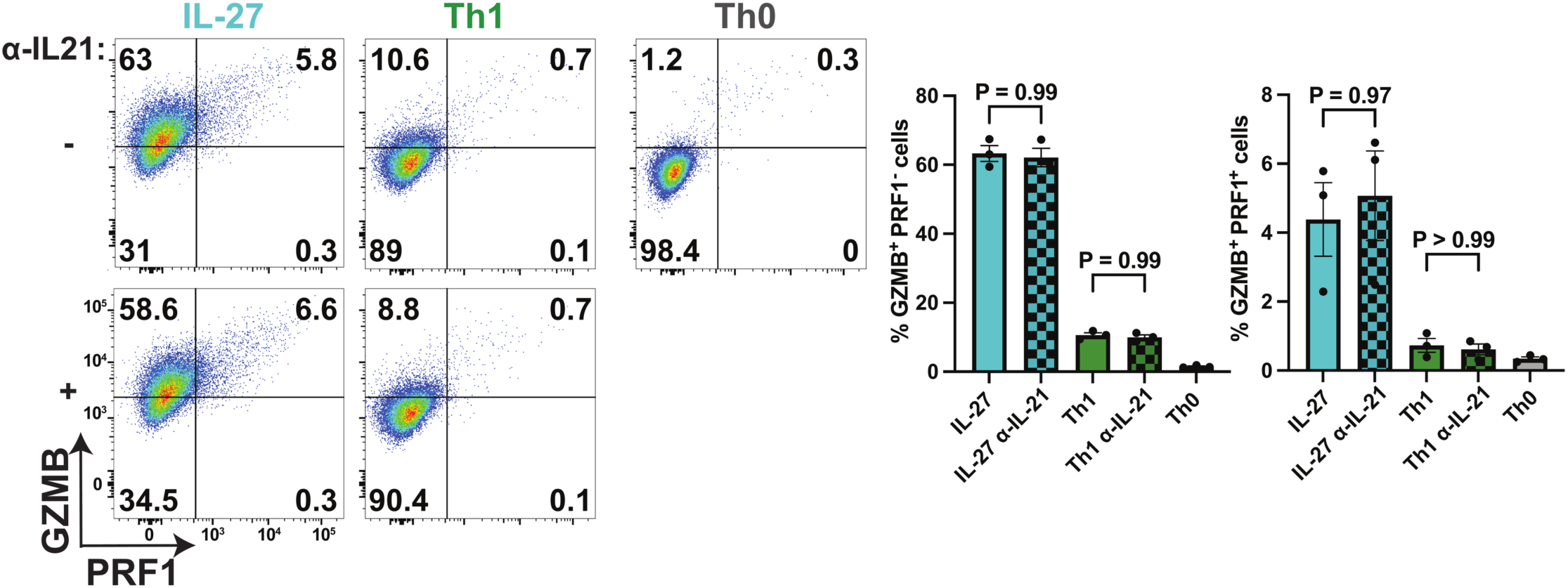
IL-27-driven cytotoxic differentiation is independent of autocrine IL-21 feedback. Naive CD4^+^ T cells from C57BL/6J mice were activated with α-CD3 and α-CD28 antibodies and polarized *in vitro* under the indicated conditions (IL-27, Th1, and Th0) for 3 days in the presence (+) or absence (–) of a neutralizing antibody targeting IL-21 (α-IL-21). **a,** Representative flow cytometry plots (left) and summary bar graphs (right) detailing the frequencies of GZMB^+^ PRF1^−^ and GZMB^+^ PRF1^+^ populations following IL-21 blockade. Data are presented as mean ± s.e.m. from n = 3 independent biological replicates. P values were determined by an ordinary one-way ANOVA with Tukey’s multiple comparisons test.

**Extended Data Fig. 6.**
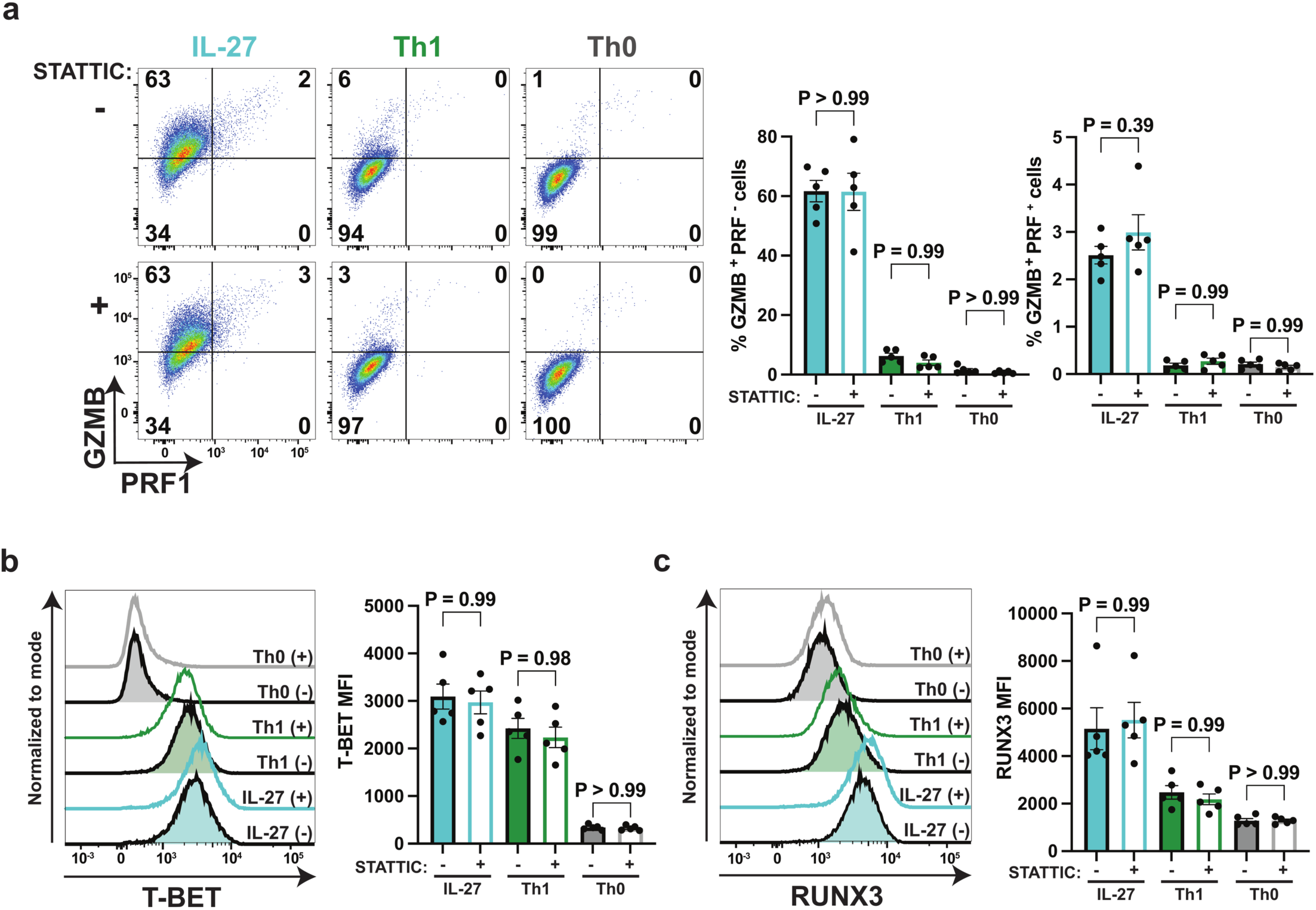
Pharmacological inhibition of STAT3 does not impair IL-27-driven cytotoxic CD4^+^ T cell differentiation. Naive CD4^+^ T cells from C57BL/6J mice were activated with α-CD3 and α-CD28 and polarized *in vitro* under the indicated conditions (IL-27, Th1, and Th0) for 3 days in the presence (+) or absence (–) of the STAT3 inhibitor STATTIC at 2 µM. **a,** Representative flow cytometry plots (left) and summary bar graphs (right) detailing the frequencies of GZMB^+^ PRF1^−^ and GZMB^+^ PRF1^+^ populations across the indicated treatments. **b,c,** Representative flow cytometry histograms (left) and summary bar graphs (right) quantifying the mean fluorescence intensity (MFI) of T-BET **(b)** and RUNX3 **(c)** across polarization conditions and treatments. Data are presented as mean ± s.e.m. from n = 5 independent biological replicates. P values were determined by an ordinary one-way ANOVA with Tukey’s multiple comparisons test.

**Extended Data Fig. 7.**
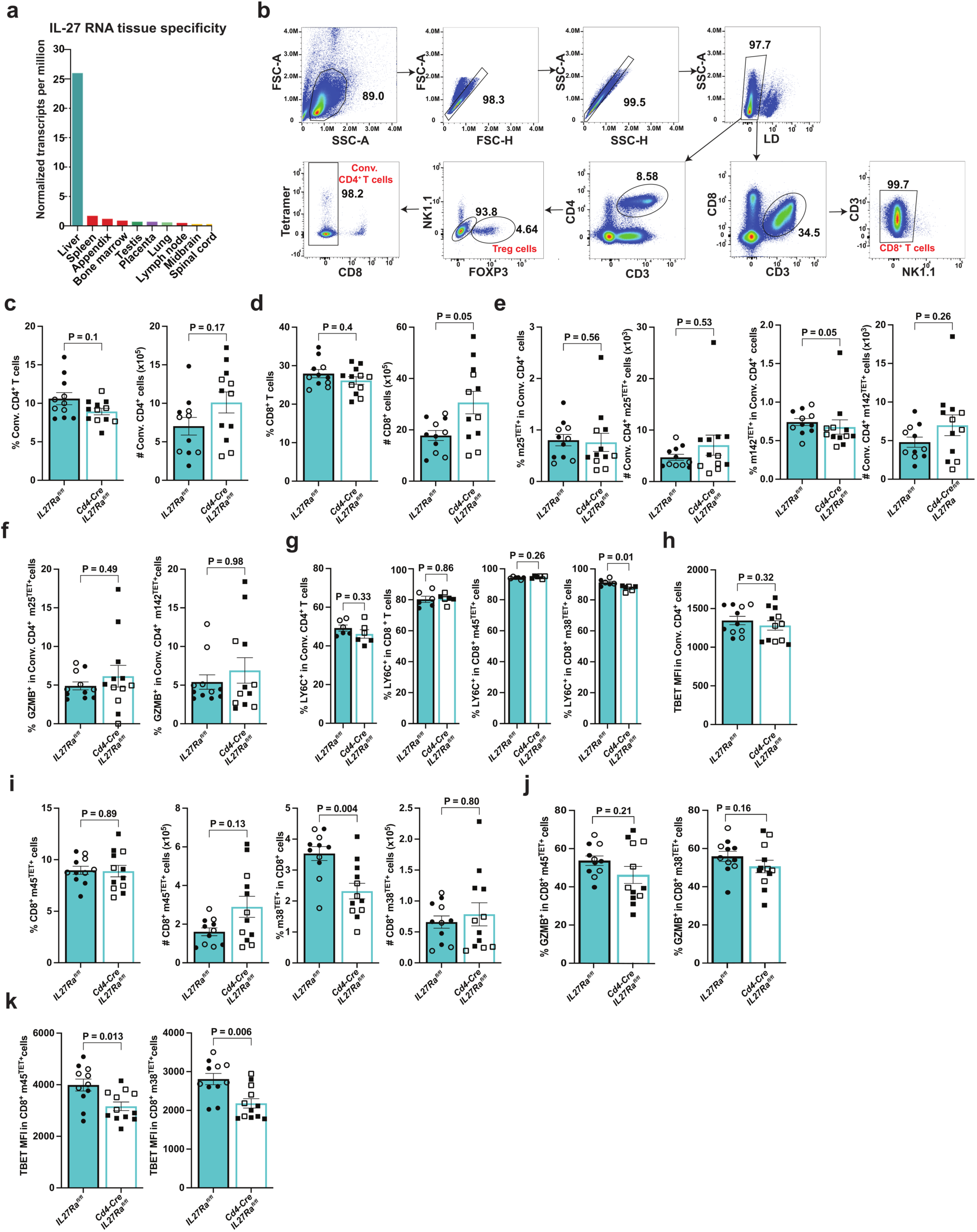
Extended phenotyping of T cell populations during MCMV infection. **a,** RNA tissue specificity data derived from the Human Protein Atlas, displaying the top 10 tissues with the highest normalized transcript expression (nTPM) for *IL27*. **b,** Gating strategy for conventional CD4^+^ T cells, Treg cells, and CD8^+^ T cells. **c–k,** *Il27ra*^fl/fl^ (control) and *Cd4-Cre Il27ra*^fl/fl^ (conditional knockout) mice were infected with 10^6^ pfu of the Smith strain of murine cytomegalovirus (MCMV) and analyzed at day 8 post-infection. Panels **c–k** show cells isolated from the liver, except panel **f**, which shows cells from the spleen. **c,d,** Summary bar graphs quantifying the frequency (left) and absolute number (right) of conventional CD4^+^ T cells **(c)** and CD8^+^ T cells **(d**). **e,** Summary bar graphs detailing the frequencies and absolute numbers of MCMV-specific m25-tetramer^+^ (m25^TET+^, left) and m142-tetramer^+^ (m142^TET+^, right) cells within the conventional CD4^+^ T cell compartment. **f,** Frequencies of GZMB^+^ cells within the m25-tetramer^+^ (m25^TET+^, left) and m142-tetramer^+^ (m142^TET+^, right) conventional CD4^+^ T cell populations isolated from the spleen. **g,** Frequencies of LY6C^+^ cells in the conventional CD4^+^ compartment, or conventional and antigen-specific CD8^+^ T cells. **h,** Mean fluorescence intensity (MFI) of T-BET in conventional CD4^+^ T cells. **i,** Percentage and absolute numbers of antigen-specific CD8+ T cells. **j,** Frequencies of GZMB^+^ cells within CD8^+^ m45-tetramer^+^ (m45^TET+^, left) and m38-tetramer^+^ (m38^TET+^, right) T cells. **k,** T-BET MFI in m45-tetramer^+^ (m45^TET+^, left) and m38-tetramer^+^ (m38^TET+^, right) CD8^+^ T cells. Data in **c–k** are presented as mean ± s.e.m. from n = 11 (*Il27ra*^fl/fl^) and n = 12 (*Cd4-Cre Il27ra*^fl/fl^) independent biological replicates (mice) except for panel **g**, where n= 6 per genotype. Solid symbols denote female mice and open symbols denote male mice. P values were determined by a two-tailed Mann-Whitney test.

## Notes

### Competing Interest Statement

The authors have declared no competing interest.

